# Visualization and quantification of coral reef soundscapes using *CoralSoundExplorer* software

**DOI:** 10.1101/2024.04.05.588225

**Authors:** Lana Minier, Jérémy Rouch, Bamdad Sabbagh, Frédéric Bertucci, Eric Parmentier, David Lecchini, Frédéric Sèbe, Nicolas Mathevon, Rémi Emonet

## Abstract

Despite hosting some of the highest concentrations of biodiversity and providing invaluable goods and services in the oceans, coral reefs are under threat from global change and other local human impacts. Changes in living ecosystems often induce changes in their acoustic characteristics, but despite recent efforts in passive acoustic monitoring of coral reefs, rapid measurement and identification of changes in their soundscapes remains a challenge. Here we present the new open-source software *CoralSoundExplorer* (https://sound-scape-explorer.github.io/docs/CSE/), which is designed to study and monitor coral reef soundscapes. *CoralSoundExplorer* uses deep learning approaches and is designed to eliminate the need to extract conventional acoustic indices. To demonstrate *CoralSoundExplorer*’s functionalities, we use and analyze a set of recordings from three coral reef sites, each with different purposes (undisturbed site, tourist site and boat site) located on the island of Bora-Bora in French Polynesia. We explain the *CoralSoundExplorer* analysis workflow, from raw sounds to ecological results, detailing and justifying each processing step. We detail the software settings, the graphical representations used for visual exploration of soundscapes and their temporal dynamics, along with the analysis methods and metrics proposed. We demonstrate that *CoralSoundExplorer* is a powerful tool for identifying disturbances affecting coral reef soundscapes, combining visualizations of the spatio-temporal distribution of sound recordings with new quantification methods to characterize soundscapes at different temporal scales.

**Author summary:** Today’s scientists are faced with the challenge of analyzing large amounts of data, such as those generated by passive acoustic monitoring of ecosystems. We have built *CoralSoundExplorer* (https://sound-scape-explorer.github.io/docs/CSE/), an efficient tool for analyzing large datasets of sound recordings, which transforms coral reef soundscape recordings into visual representations in 2D or 3D spaces. By spreading them across easy-to-explore acoustic spaces, *CoralSoundExplorer* enables the observer to quickly grasp the characteristics of soundscapes, their differences and similarities, and their organization on different temporal scales. These acoustic spaces and their temporal dynamics can be quantified, for example to account for the speed at which soundscapes change over time. In this study, we take the example of reef soundscapes from the island of Bora-Bora to illustrate the features and possibilities offered by *CoralSoundExplorer*. *CoralSoundExplorer* is open source and easy to use, even for non-specialists, thanks to an interface that requires no coding skills. We provide detailed instructions for installing and using *CoralSoundExplorer* to help users get started easily. *CoralSoundExplorer* needs to be installed on a computer to perform the analyses and calculations based on sound recordings. There is also an online interface (https://sound-scape-explorer.github.io/coral-sound-explorer/) enabling users to visualize data that have already been processed by *CoralSoundExplorer*.

## Introduction

In the current context of increasing natural habitat degradation, assessing ecosystem biodiversity and monitoring its spatio-temporal dynamics at various scales is an absolute necessity for implementing effective conservation policies, particularly for biodiversity hotspots such as tropical forests and coral reefs [1–6]. This requires rapid and reliable methods for processing large amounts of data [7]. Over the past decade, Passive Acoustic Monitoring has emerged as a crucial tool for monitoring terrestrial and marine environments, as soundscapes, at least for their biophony part, are reliable proxies for biodiversity [8–11]. By utilizing sounds as cues to the presence and activity of organisms, Passive Acoustic Monitoring provides a non-invasive, cost-effective approach to long-term monitoring of biodiversity with allowing high temporal resolution [12–15].

Coral reefs host some of the highest levels of biodiversity on the planet and provide many ecosystem services, direct or indirect benefits to several hundred million people worldwide: coastal protection, building materials, food and income from fishing or tourism [16]. However, most coral reefs are currently experiencing severe declines under the impacts of global change [17–19]. To obtain the information needed to deploy effective conservation and management policies, it is essential to have tools for spatio-temporal monitoring of the biodiversity of these reefs [20,21]. Since coral reefs are highly sonorous worlds due to the activity of numerous organisms [15,22–25], Passive Acoustic Monitoring is emerging as a particularly interesting tool for monitoring biodiversity and its dynamics [26,27]. The sounds produced by many fish species can provide information on their reproductive activity and enable changes in population abundance to be identified [28]. Sounds produced by invertebrates can also serve as indicators. For example, the density of snaps produced by shrimps is correlated to the oxygen content of the water [29]. It has also been shown that close but distinct seascapes (e.g., fringing reef, barrier reef, outer slope) are characterized by different soundscapes [26,30,31]. The soundscapes of coral reefs thus inform as to the conditions of the habitats and communities they harbor [14,26,27,32]. Moreover, they do not provide a static image but can reflect changes over time, at different temporal scales [15,33]. As a sort of multidimensional spatial and temporal picture, coral reef soundscapes are thus witnesses to biodiversity and species richness [28,34–36].

While the benefits of using Passive Acoustic Monitoring for coral reefs are commonly admitted with well-developed recording techniques at cost fairly acceptable, there remains a major challenge: developing effective analysis tools capable of apprehending large amount of data in a reasonable time and providing both qualitative and quantitative information easy to interpret [7,37]. Manual exploration of soundscape recordings by visual exploration of spectrograms and recording tracks listening is extremely time-consuming and observer-dependent [8,38,39]. The gain brought by the Passive Acoustic Monitoring approach in terms of ease of data accumulation (hydrophones are easy to install and quite inexpensive) is thus frequently diminished by an analysis process just as time-consuming and tedious as traditional survey methods (video footage, photography, and visual census).

Soundscapes are commonly described using acoustic indices [40]. In terrestrial environments, a chosen set of acoustical indices can be used to perform statistical data analysis in order to infer ecological results. Correlation studies and dimension reduction techniques can be applied to sets of acoustic indexes to explore differences between soundscapes and their evolution. From a set of indices, clustering methods can be used to perform semi and unsupervised analysis using machine learning tools. Such approaches have already been applied on coral reef soundscape [41,42]. However, there is a growing body of experimental evidence suggesting that terrestrial-derived acoustic indices are not ideal metrics to characterize underwater soundscapes [5,30,43]. More recent approaches use deep learning techniques to get a set of sound descriptors from artificial neural networks. Unlike acoustic indices, the descriptors given by a neural network are not easily intelligible but because of their efficiency in many sound analyzing tasks and their capacity to process fast on big database, deep learning approaches applied to soundscape analysis has led over the past few years to several processing workflow papers publications for a large set of bioacoustics and ecoacoustics problematics [7,9,44–50]. Among the scientific publications using artificial intelligence in the framework of ecoacoustics, the paper of Sethi et al. (2020) [51] inspired the present work. These authors proposed to analyze natural soundscapes using the pretrained VGGish neural network. From the 128 features per sound sample given by this neural network, a non-linear dimension reduction down to two dimensions was performed, allowing the visualization of the sound samples constituting the soundscapes in an acoustic space where similar sounds are mostly close to each other while different sounds are mostly distant to each other. This approach enabled visual exploration of soundscapes phenology. While the analysis described in Sethi et al (2020) [51] was appealing, it remained limited to visual inspections of soundscapes. One of our aims is to complement this approach with a quantitative method that provides reliable measures of the proximities and differences between the elements composing the soundscapes. Our other aim is to provide the user with an easy-to-use tool, where Sethi et al. (2020) [51] only offered a workflow indicating how to use a set of existing or specially developed programming language packages.

Here we present a powerful new analytical tool accompanied by an easy-to-grasp user interface, called *CoralSoundExplorer*, which uses AI deep integration technique to analyze, describe, and quantify underwater soundscapes. The workflow followed by *CoralSoundExplorer* makes use of Convoluted Neural Network (CNN) embedding and Uniform Manifold Approximation and Projection (UMAP, [52]), a dimensionality reduction technique designed to efficiently capture complex patterns and relationships in high-dimensional data. Using deep learning approaches, *CoralSoundExplorer* automatically extracts relevant acoustic features from underwater sound recordings, providing valuable information for biodiversity assessment, habitat monitoring and understanding the temporal dynamics of coral reef ecosystems to identify disturbances.

As illustrated in Fig 1, this paper is organized in a succession of sections as follows. In section I, we present the general characteristics of a coral reef soundscape and describe the recordings made on the island of Bora-Bora, which are used as sample data in this article. In Section II, we outline the principle of sound signal processing by *CoralSoundExplorer* software, the graphical representations and analyzes it enables, and the metrics that can be measured. In Section III, we show, as an illustrative example, the results obtained by *CoralSoundExplorer* with sound recordings made on the coral reef of the island of Bora-Bora. We explain how *CoralSoundExplorer* can be used to quickly and easily identify soundscapes according to the anthropogenic pressure or other perturbations they are subject to, and to capture their temporal dynamics. We describe how the analyses carried out with *CoralSoundExplorer* make it possible to account for these differences and dynamics. In section IV, we detail the methodology used by *CoralSoundExplorer* to analyze the recordings and measure the metrics. Although the main methodological aspects are presented in the previous sections, the details are only covered in this section to make the document easier to read. Section V examines the main achievements and limitations of *CoralSoundExplorer*, and presents our recommendations for exploring reef recordings with our software. Section VI is a guide to install and use *CoralSoundExplorer*.

**Fig 1.**
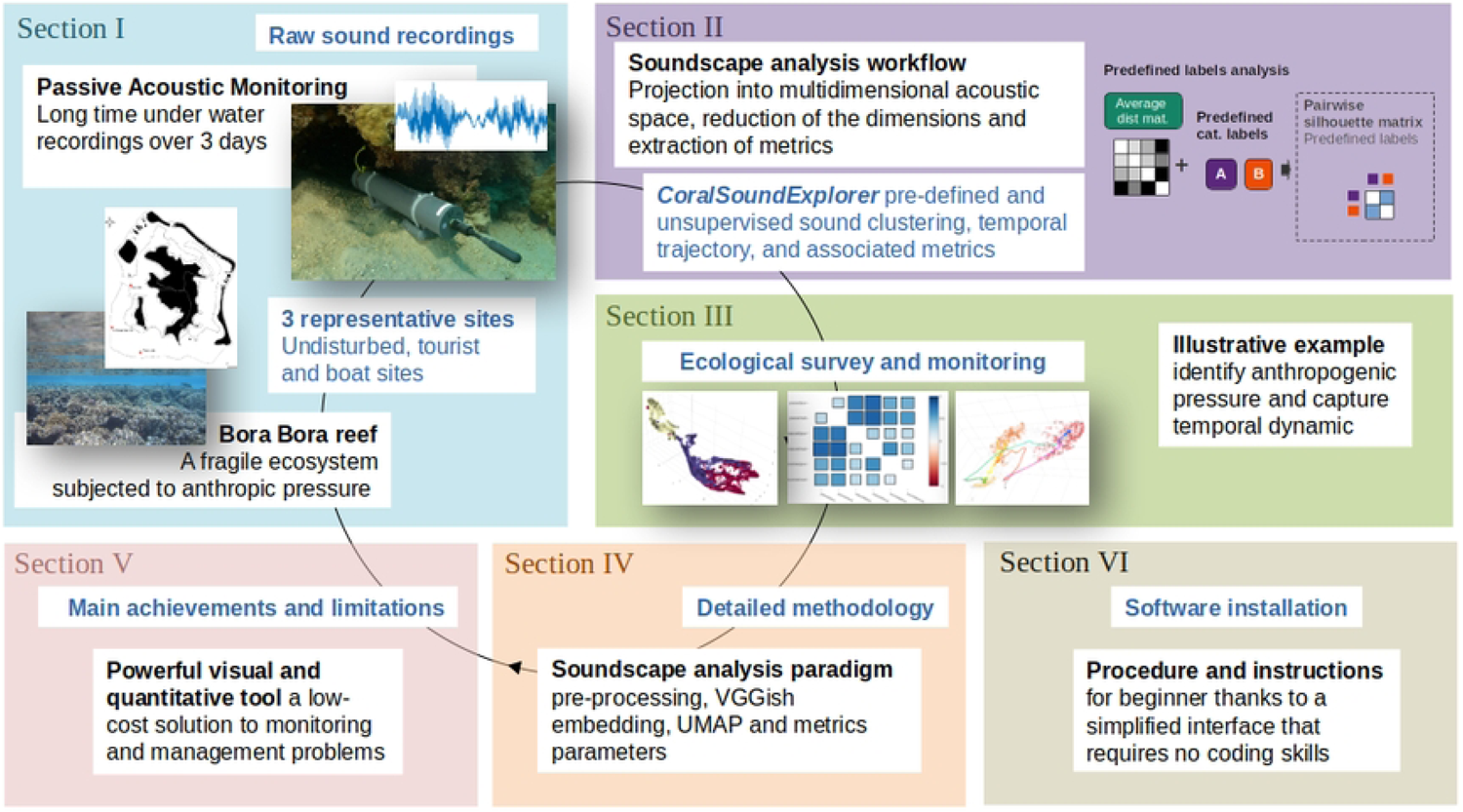
Study framework. Section I: General characteristics of a coral reef soundscape and presentation of the recordings made on the island of Bora-Bora that serve as example data in the present study. Section II: Presentation of *CoralSoundExplorer* software (analysis workflow, graphical output and measurable metrics). Section III: Results obtained with *CoralSoundExplorer* from the sample recordings made on Bora-Bora. Section IV: Detailed methodology. Section V: Main achievements and limitations. Section VI: Instructions for software installation.

## I. The soundscapes of coral reefs and the example of Bora-Bora Island

### 1. What is a coral reef soundscape?

Like all aerial or underwater soundscapes, those of coral reefs are the result of an interweaving of sounds produced by animals (biophony - Fig 2A & Fig 2B), sounds from human activities such as shipping (anthropophony - Fig 2C) and sounds emanating from geophysical processes such as waves breaking on the reefs (geophony - Fig 2D). As coral reefs are among the world’s richest ecosystems in terms of biodiversity [53,54], their soundscapes are among the most complex of aquatic realms. Fish make sounds when they eat or swim and most of them produce sounds to communicate, notably during agonistic interactions, in response to threats such as the presence of predators, or during courtship and spawning [23] (Fig 2B). While up to half of the fish families may have sonorous species, benthic invertebrates also play a major role in coral reef biophony (e.g., 55–57). Snapping shrimps, common on coral reefs, are very noisy [55]. Bivalves, clawed and spiny lobsters and burrow-dwelling mantis shrimps also produce sounds [56]. Sea urchins when moving or grazing also contribute to biophony [24,58]. As coral reefs vary according to depth, hydrodynamic conditions and other parameters, there is a wide diversity of animal species assemblages and associated habitats [30]. This ecological diversity drives a diversity of soundscapes that reflect the properties of the coral reef ecosystem, in its spatial and temporal components, and are thus witnesses to the diversity of fish and other animals, as well as coral cover [14,26,27].

**Fig 2:**
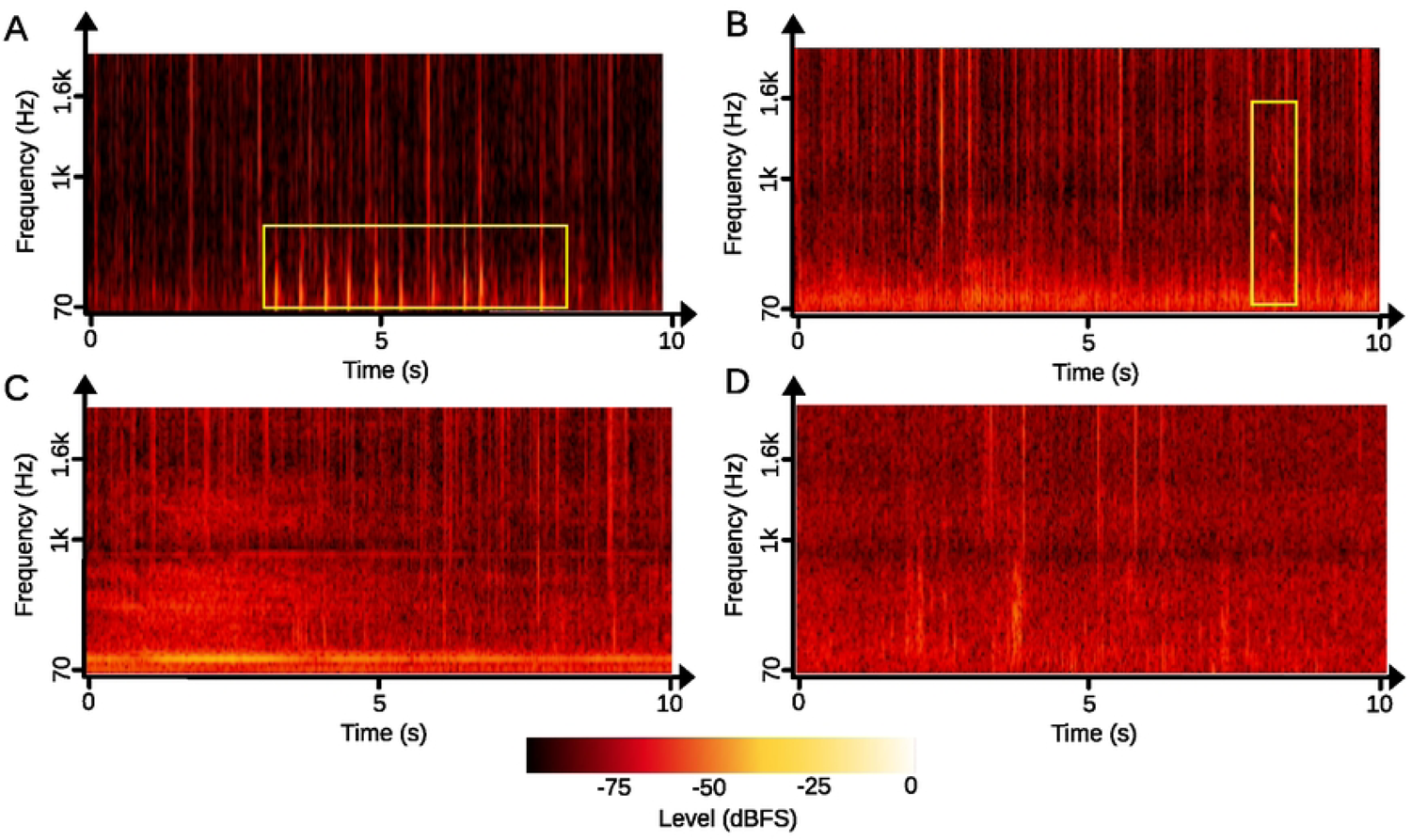
Soundscapes diversity in a coral reef. (A) Spectrogram of a reef soundscape recorded during the day. Fish sounds (yellow square) are mostly concentrated at low frequencies (< 1 kHz). (B) Spectrogram of a reef soundscape recorded at night showing an overall increase of sounds levels mostly due to an increase in the number of snapping shrimp sounds. An unidentified fish sound is also present (yellow square). (C) Spectrogram showing the broadband noise of a passing boat, with higher intensity during the first 5 seconds. (D) Spectrogram showing broadband masking effect caused by rain. Spectrograms were plotted with *CoralSoundExplorer*, using FFT window size = 2048 samples, sampling frequency = 44100 Hz.

### 2. The example of Bora-Bora: Reefs under heterogeneous anthropic pressure

Bora-Bora, a tropical volcanic island in French Polynesia (16°29’ S, 151°44’ W), is an excellent example of several coral reefs habitats: fringing reef, channel, barrier reef, mangrove, *etc.* [59]. Bora-Bora is surrounded by a coral reef of 70 km² which is home to a high diversity of fish, invertebrates and other marine organisms [3]. Despite being a renowned international tourist destination with luxurious resorts and environmental commitment, it faces anthropogenic pressures, mainly from intense tourism [3,60,61]. However, such tourism success is not without impact. It generates significant maritime traffic, inevitably accompanied by noise pollution caused by the intensive use of internal combustion engine-powered boats. To regulate the presence of tourist activities in the lagoon, 14 ecotourism sites have recently (2019) been identified by the town council and the tourism committee [3,62]. Despite these measures, some sites on Bora-bora remain heavily impacted by tourist activities, not to mention the fishing activities that meet the food needs of a large part of the population [3,63]. To give an idea of the scale of nautical activity, 711 boats for personal use such as sailing, leisure and fishing are based on Bora-Bora [64].

### 3. The Bora-Bora coral reef dataset

We selected three sites in the south-western part of Bora-Bora lagoon (Fig 3): a tourist site (16°32’11.543” S, 151°43’30.575” W) used for snorkeling activities with healthy corals and a high fish species richness (max. 2 m depth), an undisturbed site (16°31’46.956” S, 151°47’19.823” W) where very few boats can access due to a high coral cover and which has no tourist activity; and a boat site where boat traffic is particularly intense in a deep sandy channel (16°30’7.416” S, 151°46’5.448” W; [3,15]. We carried out three passive acoustic recording sessions in November and December 2022. The duration of the recordings was deliberately limited to enable parallel manual analysis of the sound recordings and comparison with the results obtained with *CoralSoundExplorer* software. This dataset was therefore suitable for checking the results obtained with *CoralSoundExplorer*.

**Fig 3:**
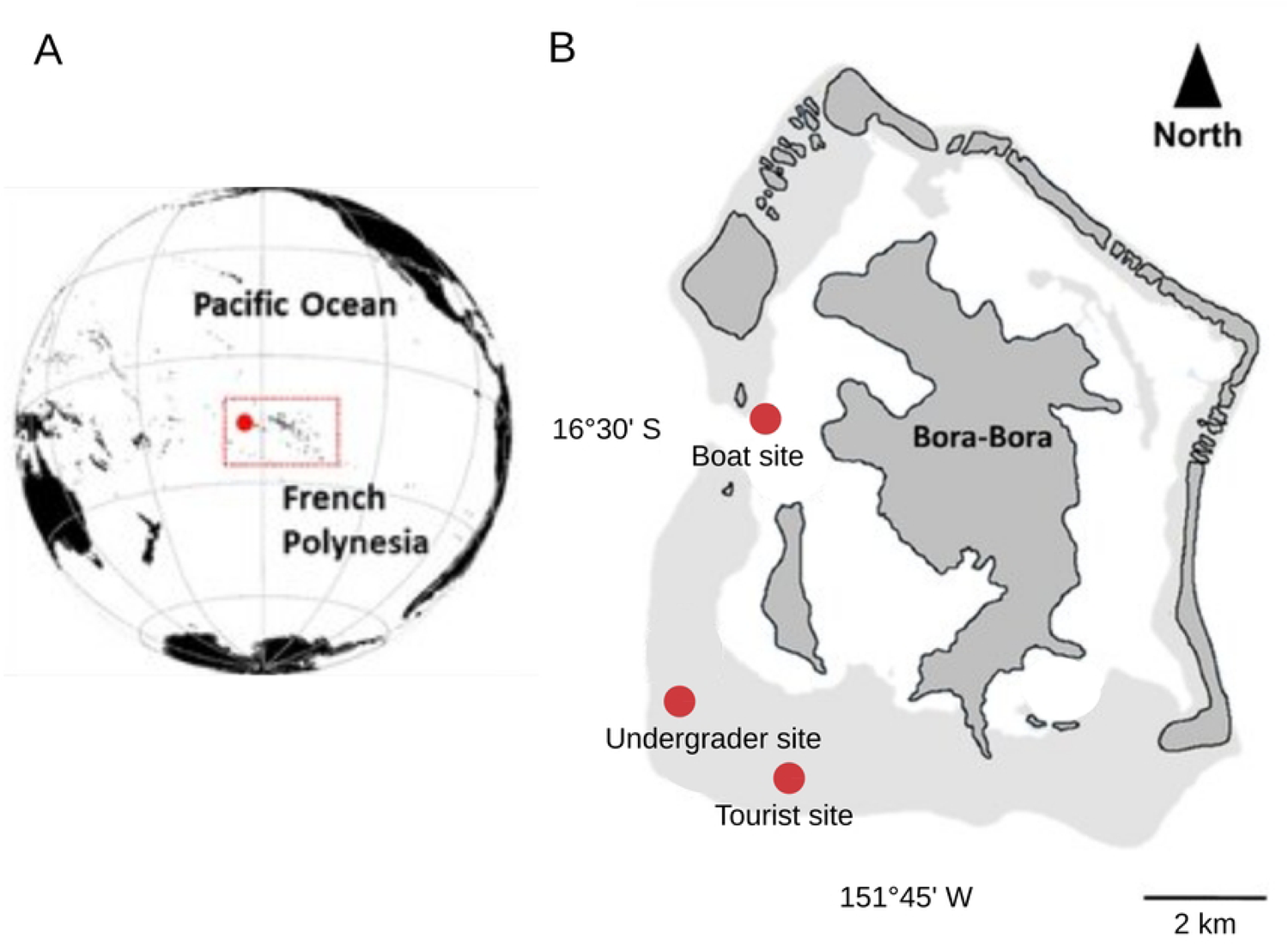
Location of recording sites on Bora-Bora Island. (A) World map showing the location of the French Polynesia archipelago (red square) and the world distribution of coral reefs area (blue square). (B) Map of the Society islands in French Polynesia showing the location of Bora Bora (red square). (C) Map of Bora-Bora showing the three recording sites (boat site, undisturbed site, and tourist site). In black: land areas; black line encircles the reef area.

On each of the three sites and for three different days (November 16, November 22 and December 12), we deployed simultaneously SNAP long-term autonomous acoustic recorders (Loggerhead Instruments; Sarasota, FL, USA) equipped with a HTI-96-Min hydrophone (sensitivity 169.9 dB, 170.0 dB, and 170.1 dB re 1 V at a sound pressure of 1 µPa; flat frequency response from 2 Hz to 30 kHz). The recorder positioned at each of the three sites recorded the soundscape over 24 h, from 12:05 p.m. to 12:05 p.m. (included) the next day, with a recording cycle of 1 min every 10 min (sampling frequency 44.1 kHz; 16-bit resolution). On all three sampling dates, the three recorders were systematically positioned in exactly the same spot, on a sandy substrate beneath a coral head, at depths of 1 m, 2 m and 5 m for the tourist, undisturbed and boat sites respectively.

The original recordings were high-pass filtered with an 8-order zero-phase Butterworth filter set at a cut-off frequency of 70 Hz. The soundscape analyses were carried out in the low-frequency band of interest, between 70 Hz and 2 kHz where most fish sounds and anthropogenic noises are found [65,66]. All recordings were labeled according to location (undisturbed, tourist or boat sites), time of day (daytime or nighttime, depending on the exact time of sunrise and sunset), and date (replicate 1 for November 16, replicate 2 for November 22, and replicate 3 for December 12). A composite label composed of all three previous labels has been created to enable simultaneous and combined analysis of these three factors. This combination of initial labels produced 18 composite labels (3 x 2 x 3). We also listened to all these recordings, associating them with metadata such as the presence of rain, fish sounds or motorboat noises.

## II. *CoralSoundExplorer* software: Analysis workflow, graphical output and measurable metrics

The soundscapes analysis workflow handled by *CoralSoundExplorer* consists of three main steps: the projection of sounds into a multidimensional acoustic space, the reduction of the dimensions of this space so that it can be visualized in 2D or 3D, and the extraction of metrics characterizing the organization of the sounds composing the soundscapes. This part II briefly presents the *CoralSoundExplorer* analysis workflow and the results that can be drawn from it. Technical details, analysis parameters and their rationale are presented in detail in Part IV.

### 1. From original sound signals to acoustic space

*CoralSoundExplorer*’s workflow is illustrated in Fig 4, from original sound signals to graphical representations and possible calculations. The first part of the workflow (Fig 4A) is partly inspired by previously published work [51]. The original one-minute raw recordings are first divided into one-second signals. These one-second signals are then processed using a Short-Time Fourier Transform (STFT) leading to a two-dimensional representation of sounds (time/frequency representation). Each of these 2D sound representations, one per raw signal, is then presented to the input of a pre-trained Convolutional Neural Network (CNN) called VGGish [67]. The output of this CNN is a projection into a high-dimensional acoustic space (128 dimensions), called VGGish embedding, where each second of recording is assigned 128 dimensions related to its acoustical characteristics. Once the entire dataset has been projected into the VGGish embedding, a process of deleting unnecessary dimensions (i.e., dimensions containing no information) is carried out, followed by a normalization process (robust scaling) on each remaining dimension. A temporal grouping of the one-second signal projections is then performed by calculating the average position in the acoustic space for every 15 consecutive seconds of a recording, without overlap. This averaging results in 4 points per one-minute recording (instead of the initial 60 points per minute). This 15-second average was chosen because the recorded soundscapes of the Bora Bora coral reef are relatively homogeneous and representative on this time scale. The 15-second duration is short enough in relation to the sounds emitted by the boats to capture these singularities and long enough to avoid the strong influence of isolated, shorter duration sound events. However, the *CoralSoundExplorer* user is free to set this time averaging to any desired value, the minimum being one second. Because we have recorded for 24 hours from 12:05 p.m. to 12:05 p.m. (included) the next day, on three sites for three days, the entire Bora-Bora dataset was composed of 5220 samples (4×3×(6×24+1)×3). After the deletion of unnecessary VGGish dimensions considering the bora-bora dataset (see: section IV Detailed methodology: Signal pre-processing and VGGish embedding), each of these 5220 points was characterized by 119 dimensions. A dimension reduction process is then carried out using an implementation of the UMAP algorithm to obtain a sufficiently low number of dimensions (3 dimensions) to allow visualization of the points in an acoustic space, as well as the calculation of sound organization metrics. As the UMAP transformation process is not deterministic, there are differences in point positions between two different UMAP projections. Metrics calculated directly from these projections or from point distance matrices are also affected by this stochasticity. To mitigate these random effects of the UMAP algorithm and reduce the uncertainties of the analysis metrics, the UMAP dimension reduction process is run 100 times from the same VGGish embedding. From these 100 3D UMAPs, an average distance matrix is calculated. A 2D and/or 3D UMAP transformation is also calculated from the VGGish embedding to enable visualization of the results.

**Fig 4.**
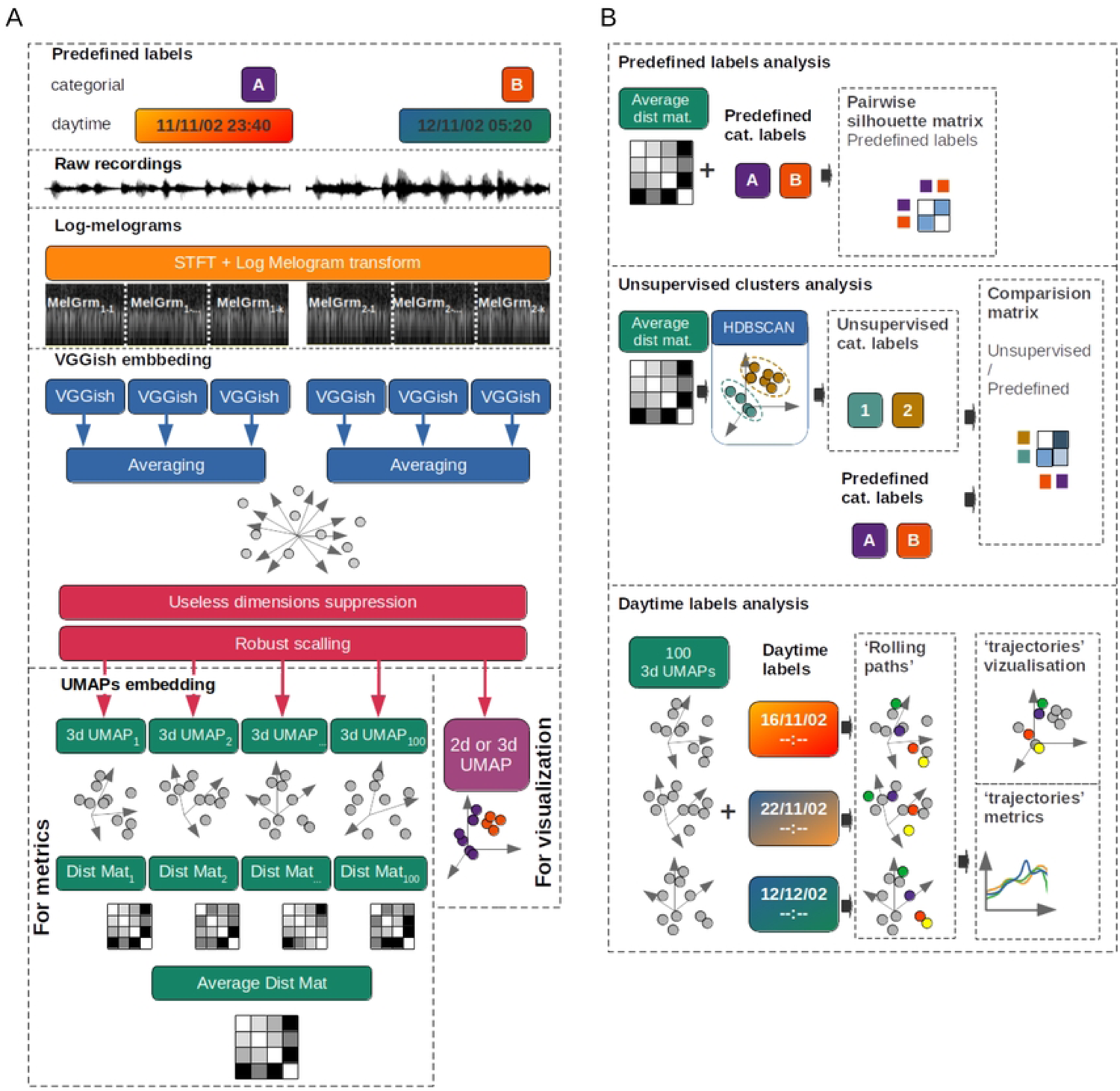
*CoralSoundExplorer* workflow. (A) **From sound recordings to graphical representations and distance matrices.** Raw sound recordings, associated with predefined discrete (categorical) or continuous labels (i.e., Handcraft labels, e.g., recording locations and times), are cut into one-second extracts and transformed into log-melograms (MelGrm). Each log-melogram is processed by a convolutional neural network (VGGish). The neural network assigns 128 descriptors to each 1-second sequence. These one-second samples are then aggregated according to the chosen integration time (15 seconds in the present study) and their 128 descriptors are averaged over this duration. Dimensions containing no information (i.e., descriptors equal to 0 or constant for the whole dataset) are eliminated. This step is followed by a normalization process on each remaining dimension (robust scaling). The population of 15-second aggregated descriptors is projected into a two- or three-dimensional space using the UMAP method. The UMAP operation is repeated 100 times. For each iteration, the distances between each pair of sounds in the acoustic space are calculated to establish distance matrices (Dist Mat_1_ ≤ n ≤ 100), which are then averaged (Average Dist Mat). (B) **Analyses based on UMAP projections. Top panel:** Evaluation of the suitability of predefined labels to describe the organization of the acoustic data. This evaluation can be made visually from the UMAP projection in the acoustic space, where each sound is identified by a colored dot according to a predefined label. It is also quantified from the average distance matrix by calculating silhouette indices (Pairwise Silhouette index). **Middle panel:** Unsupervised sound clustering. The HDBSCAN algorithm identifies sound clusters based on their proximity in the average distance matrix. These clusters are associated with so-called unsupervised labels (Unsupervised cat. Labels) that identify sounds in acoustic space. The unsupervised clusters can be visualized in a 2/3D projection (visual UMAP) and compared with clusters derived from predefined labels (Predefined cat. Labels) to assess the correspondence between the two categories of labels (Unsupervised/Predefined comparison matrix). **Bottom panel:** The temporal trajectories of soundscapes in soundspace (Rolling path) are calculated and plotted based on the position of UMAP points relative to the starting position (recording start time). This calculation is averaged over 100 UMAPs (see text for details).

### 4. Analysis 1: Assessing the suitability of predefined labels for describing data organization

The three types of analysis carried out based on UMAP projections are illustrated in Fig 4B. The corresponding *CoralSoundExplorer’s* display features are illustrated in Fig 5, 6 & 7.

**Fig 5:**
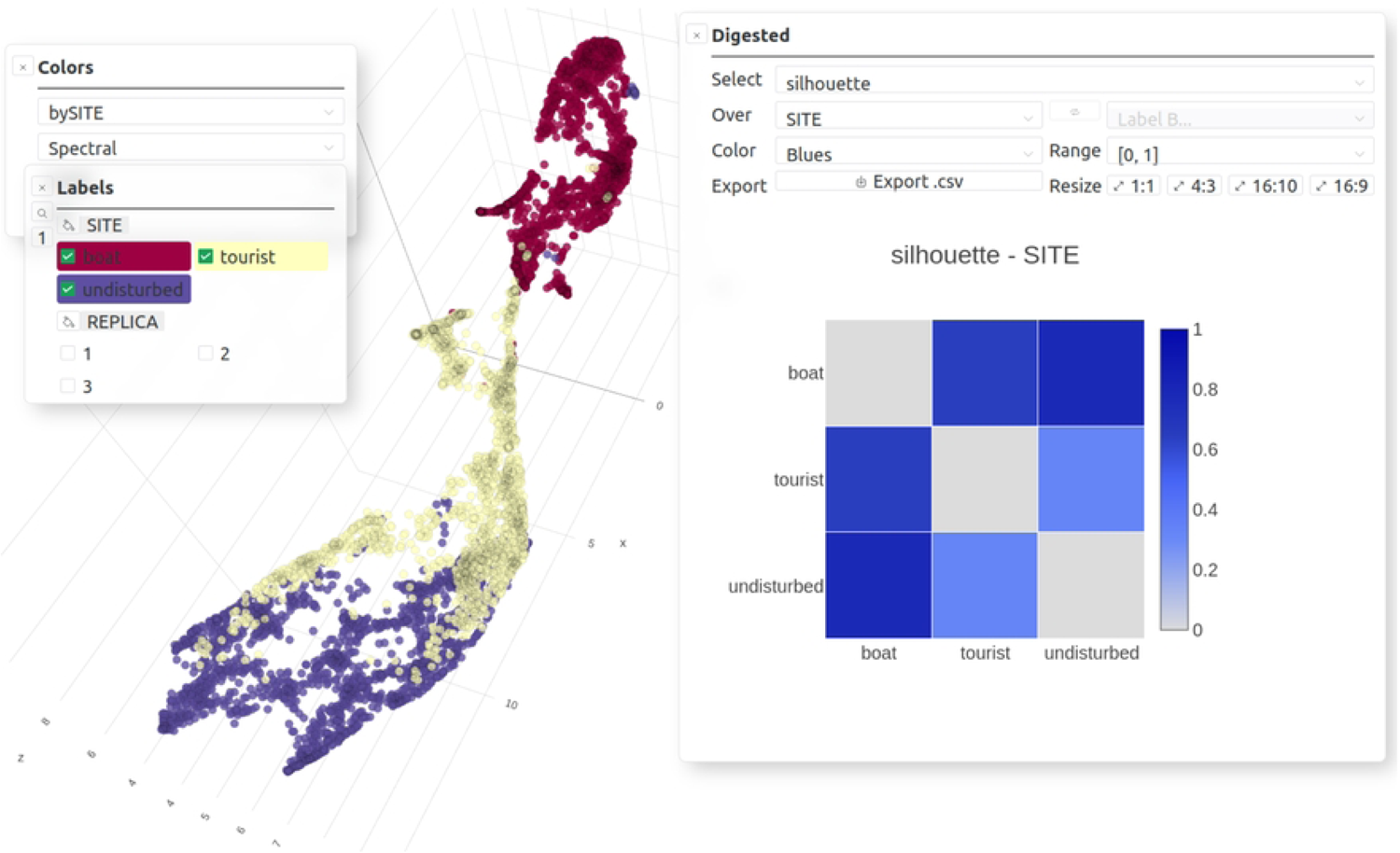
Analysis offered by *CoralSoundExplorer*: Assessing the suitability of predefined labels for describing data organization. Graphical display possibilities: Color points (sounds) according to the values of a categorical variable. Pairwise silhouette indices matrix for one categorical variable. Export possibilities: UMAP plot (.png, .svg), Pairwise silhouette indices matrix (.csv) and its plot (.png, .svg).

**Fig 6:**
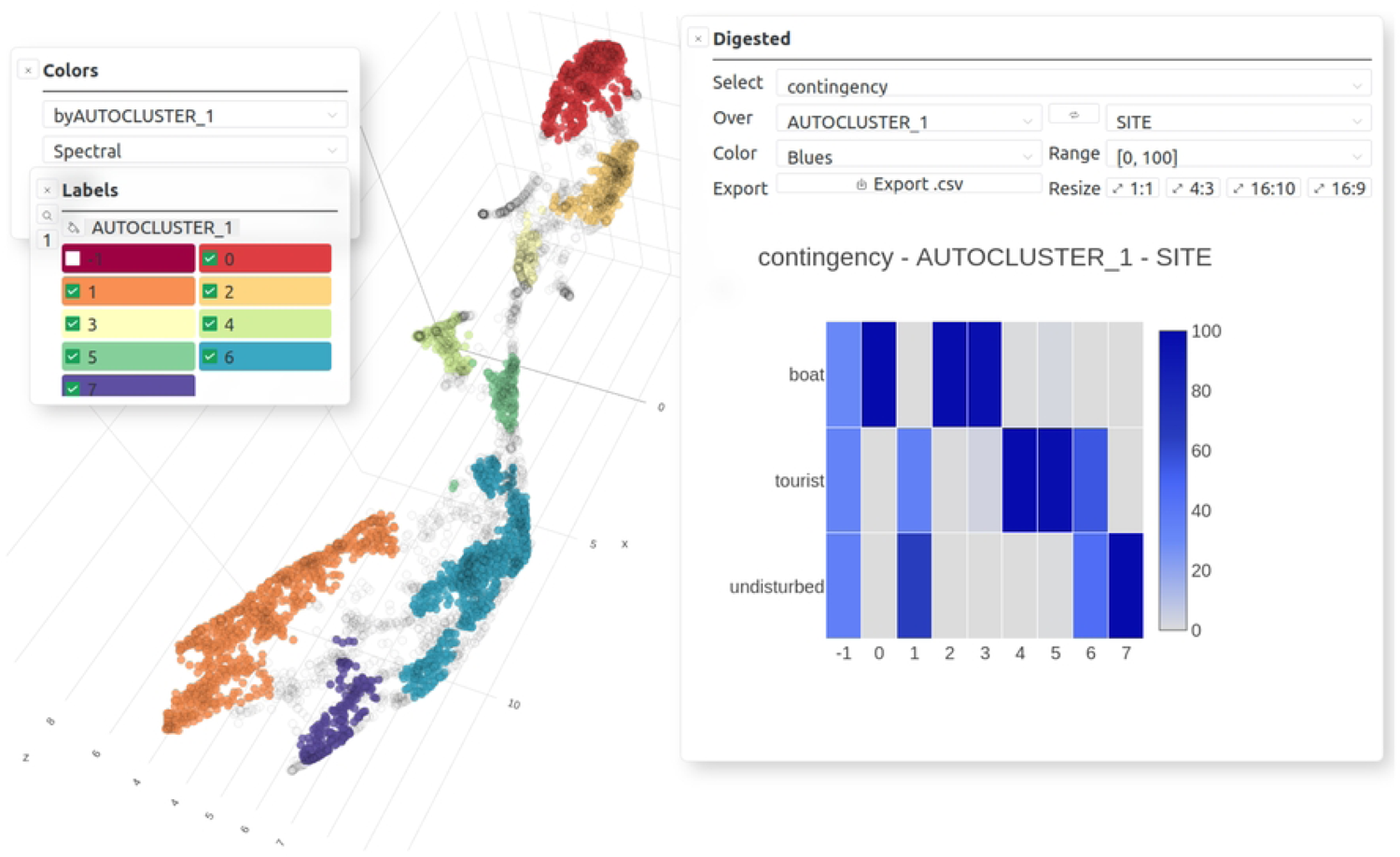
Analysis offered by *CoralSoundExplorer*: Unsupervised sound clustering. Graphical display possibilities: Color points (sounds) according to the values of unsupervised clusters id, Contingency matrix (i.e., percentage of each modality of one categorical variable within each modality of another categorical variable). Export possibilities: UMAP plot (.png, .svg), contingency matrix (.csv) and its plot (.png, .svg).

**Fig 7:**
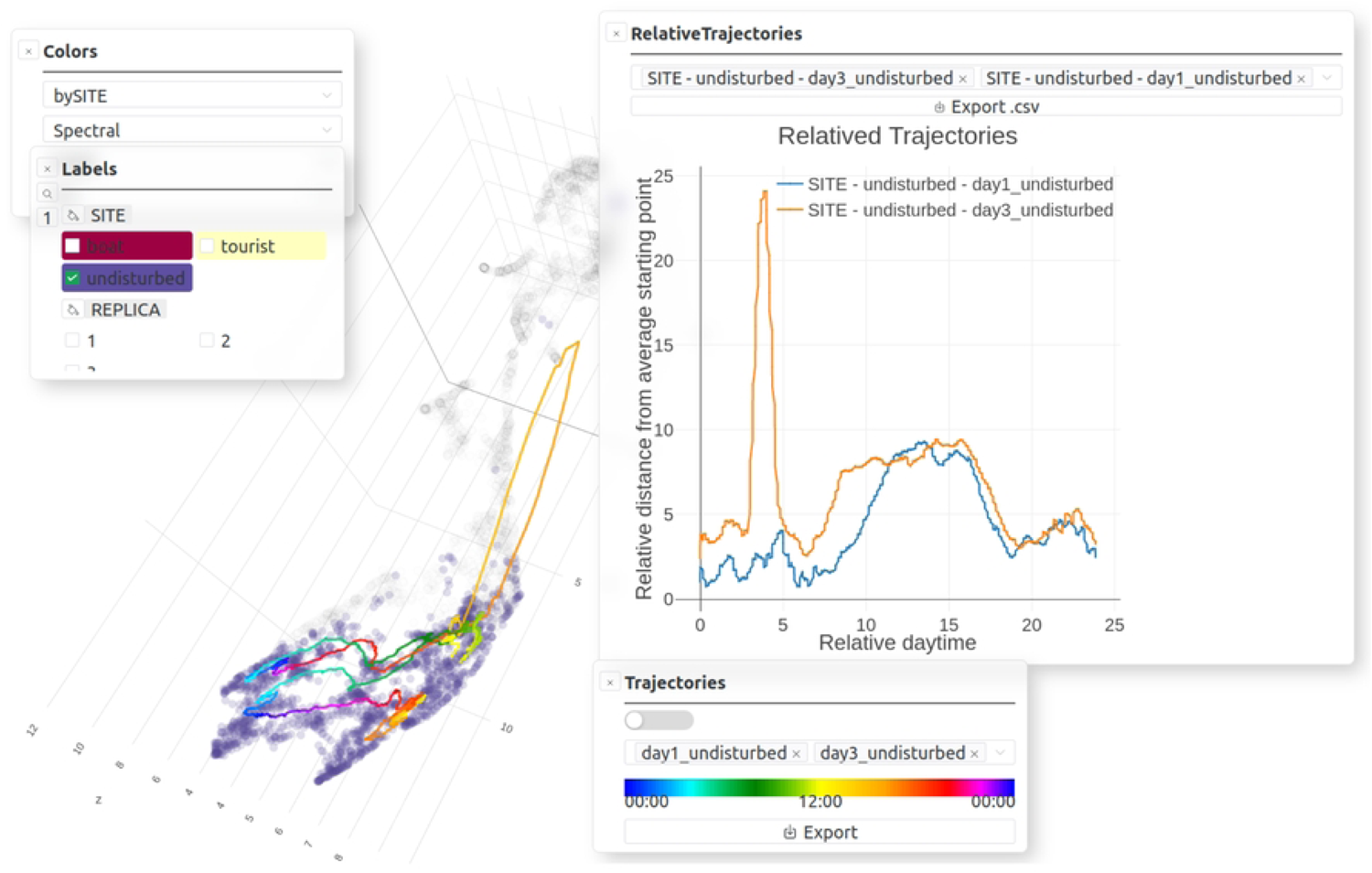
Analysis offered by *CoralSoundExplorer*: Temporal trajectory of soundscapes. Graphical display possibilities: plot predefined time trajectories, plot “Relative distances [from average starting point]”. Export possibilities: UMAP plot (.png, .svg), “Relative distances [from average starting point]” (.csv) and their plots (.png, .svg).

In the first analysis (Predefined labels analysis; Fig 4B top & Fig 5), we assess the extent to which sounds that all share the same predefined label (e.g., having all been recorded at the same location or all on the same day or same day period) effectively form an identifiable and distinct group from sounds with other labels. The assessment of this ability of predefined labels to form distinct acoustic clusters is based on the calculation of the silhouette index [68]. This index is used here to quantify the overlap of positions occupied in acoustic space by sounds bearing two different labels. The silhouette index gives an indication of the dissimilarity of sounds belonging to two labels. When the silhouette index equals 1, sounds defined by the two labels from two distinct groups in acoustic space. When the silhouette index is equal to 0, sounds belonging to both labels form undifferentiated groups, meaning that the labels assigned to the sounds do not translate into differences in position in acoustic space. The points representing the sounds of the two labels then occupy the same region of acoustic space. For a given categorical label, *CoralSoundExplorer* automatically calculates the silhouette indices between the sounds considering each of the label’s categorical values in pairs. The results are displayed as a matrix of colored squares, the color indicating the index value. This allows visual exploration of how sounds share acoustic space according to the predefined labels. The numerical values of the matrix are exportable as a table in a CSV file.

### 5. Analysis 2: Unsupervised sound clustering

In this second analysis (Fig 4B middle panel & Fig 6), *CoralSoundExplorer* seeks to identify sound clusters in an unsupervised way based solely on their analysis by the neural network and the 100 UMAPs average distance matrix, without prior reference to the labels previously assigned to the sounds. This search for clusters is made using the HDBSCAN clustering algorithm [69]. This algorithm identifies groups of sounds (unsupervised clusters) in the acoustic space within which the sounds are similar to and different from those of other groups. This algorithm has the advantage of working without prior knowledge of the number of clusters. It can also exclude some sounds from any cluster if it judges them to be too isolated in the acoustic space. The result of this analysis is to assign each sound in the acoustic space the label of the cluster to which it belongs or, where appropriate, to indicate that a particular sound does not belong to any of the clusters. This result can be directly visualized on the UMAP acoustic space representation in two or three dimensions. The points representing the sounds are then colored according to their cluster membership. These unsupervised clusters, automatically defined by the HDBSCAN algorithm, can then be compared with the clusters formed by the sounds bearing labels previously defined, to identify the proportion of them making up each of the unsupervised clusters. This sub-analysis is presented in the form of a matrix, with the rows corresponding to the labels of the unsupervised clusters and the columns to the categorial predefined labels. The values in this matrix correspond to the percentage of the number of sounds carrying the predefined label that belong to the different clusters defined in an unsupervised way. To allow visual assessment of these values, *CoralSoundExplorer* offers a graphical representation with colored squares, whose shades of color are proportional to the matrix values. HDBSCAN’s computation parameters are preset by default in the software. They can be adjusted to obtain different levels of analysis refinement, leading to different sizes and number of unsupervised clusters. Two different HDBSCAN settings were used in this study, which differed in the final classification process used.

### 6. Analysis 3: Temporal trajectory of soundscapes

The third analysis tracks the evolution of the soundscape over time (Fig 4B bottom panel & Fig 7). It provides a mapping visualization and associates a metric to quantify the evolution of sounds over time. Temporal monitoring of the soundscape involves defining the sub-regions of acoustic space explored by sounds over a given period and for a given recording location. This creates ‘paths’ in the acoustic space along time points. These paths are computed from the UMAP projection in 2 or 3 dimensions. They are plotted graphically in the acoustic space defined by the UMAP, to visualize the phenology of the soundscape. These trajectories are colored according to the time of day. By comparing the temporal paths of the same recording location between different days, these paths make it easy to detect outliers in the temporal trajectories of the soundscape that may indicate changes in the ecological situation (e.g., between days). It should be noted that the calculations enabling this visualization of temporal paths are originally based on a single UMAP projection. They are therefore subject to UMAP stochasticity. To give a stable metric of the evolution of the soundscape over time relative to each recording site, *CoralSoundExplorer* also makes use of the several UMAPs (here 100, see part 4: Detailed Methodology). The measure is the distance from the average starting point of each site’s paths. Because this distance in UMAPs is not absolute, it is defined relative to the average distance between its reference point (starting point) and its 100 nearest neighbors in the UMAP. For a given site, day and time, the median value of these distances (over the 100 UMAPs) is retained. These median distances are then plotted against time of day, for a given site and a given replicate. In this way, it is possible to appreciate phenology in a different way from that possible directly on the UMAP visualization. All these relative trajectories start and often end with a value around one indicating a cyclic phenology of soundscapes during the whole day/replicate. During the day, the relative trajectories can diverge, indicating a change of soundscape compared to the one present during their starting point.

### 7. Other display features and data exploration tools provided by CoralSoundExplorer’s user interface

*CoralSoundExplorer* is a tool for exploring soundscapes, combining a graphical interface for visualizing the distribution of recorded sounds in a 2- or 3-dimensional acoustic space, with functions for quantifying this distribution. *CoralSoundExplorer*’s graphical interface includes interesting possibilities for exploring a sound database. *CoralSoundExplorer* offers the option of coloring points (corresponding to the different sounds) directly from the graphical interface, according to the values taken by a predefined categorical variable or by the cluster obtained after unsupervised cluster identification. It is also possible to hide sounds in the projection space according to the value(s) of another or the same categorical variable (Fig 8A). This allows the user to quickly visualize the organization of recorded sounds according to the attribute value of a categorical variable, and to visualize possible interactions between two or more categorical variables.

**Fig 8.**
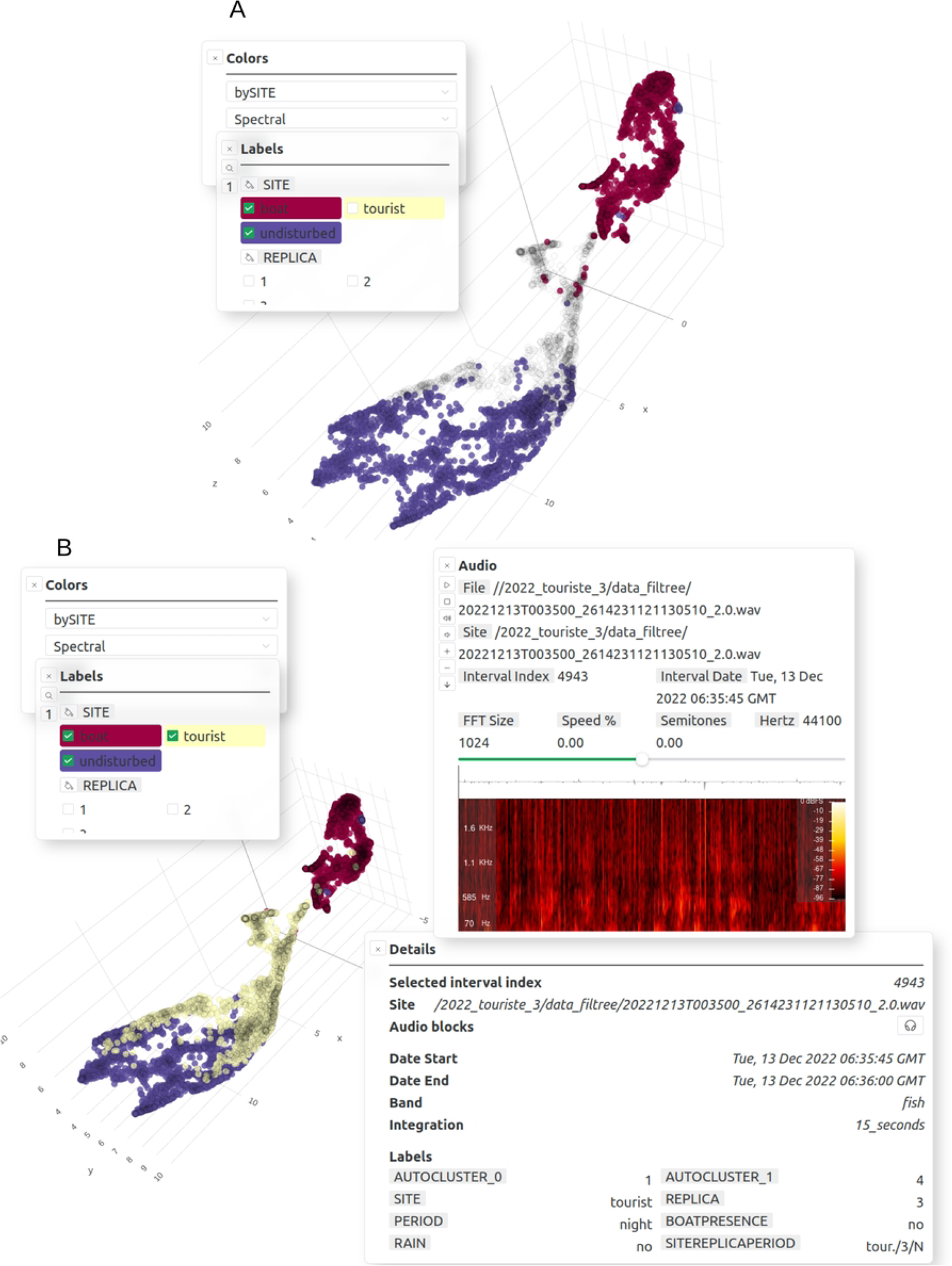
Other display facilities and data exploration tools provided by *CoralSoundExplorer* 1/2. (A) Display features: color points (sounds) according to the values of a categorical variable, highlight points according to the values of a categorical variable (the same or another). Export possibilities: UMAP plot (.png, .svg). (B) Display features: sounds information and their spectrograms after clicking on the corresponding point. An audio player is provided to listen to the selected sound. Export possibilities: sound (.wav).

Each sound sample corresponding to a point in the UMAP 2D or 3D acoustic space can be played by clicking on it (Fig 8B). The spectrogram of this signal is also displayed. The size of the spectrogram analysis window can be adjusted, the overall sound level increased or decreased, as well as the playback speed. This makes it possible to inspect any sound signal in the acoustic space on demand (by listening and viewing the spectrogram), for example to understand what in the sounds’ spectrotemporal structure may have influenced *CoralSoundExplorer*’s process of grouping points.

In addition to the temporal trajectory of the soundscape analysis, it is possible to obtain a quick overview of the phenology of sound samples using a continuous color scale of points according to the corresponding start time of the sample recording. In order to focus on the phenology of soundscapes defined by one or more categorical variable values, this functionality can be combined with the ability to color only the sound projection points in the projection space corresponding to one or more categorical variable values (Fig 9A).

**Fig 9.**
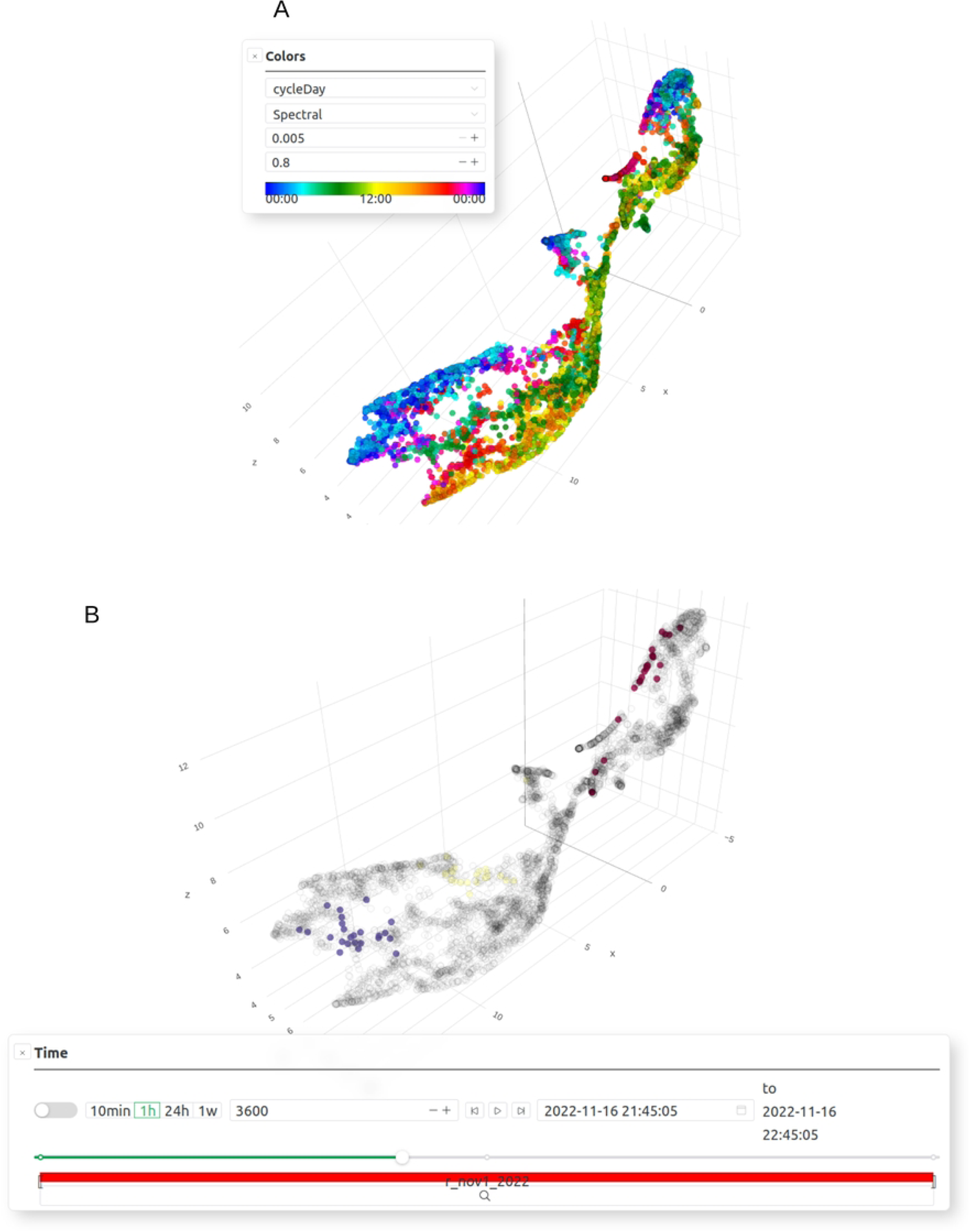
Other display facilities and data exploration tools provided by *CoralSoundExplorer* 2/2. (A) Display features: color points (sounds) using a continuous scale according to the starting time of each point (starting time of the recording). Export possibilities: UMAP plot (.png, .svg). (B) Color only points corresponding to sounds recorded in a specified time interval. Export possibilities: UMAP plot (.png, .svg).

On the UMAP visual projection, *CoralSoundExplorer* also offers the option of highlighting (or color) only those points corresponding to recordings made within a specified time interval, giving the date and time of the start of this interval and its duration (Fig 9B).

## III. *CoralSoundExplorer* software: Results obtained with the Bora-Bora dataset

The Bora-Bora dataset illustrates the main objectives of *CoralSoundExplorer*: 1) visually explore a bank of sound recordings bearing labels (recording locations, date, time) and quantify the acoustic proximity of recordings as a function of these labels, 2) explore a bank of sound recordings without using labels a priori by identifying acoustic clusters, then seek to explain these acoustic clusters as a function of labels, 3) visualize and quantify the temporal dynamics of soundscapes, enabling the detection of disturbing events, and compare the temporal dynamics of different soundscapes.

### 1. Spatio-temporal distribution of sounds with labels

Fig 10 shows the projection in 3D acoustic space of the Bora-Bora recordings processed by *CoralSoundExplorer*. This figure illustrates the possibility of coloring the recordings’ projections according to each of the predefined labels. Here, the integration time chosen is 15 seconds. Each point in the cloud therefore represents 15 consecutive seconds of recording. The set of points forming the cloud represents all the recordings in the Bora-Bora dataset.

**Fig 10.**
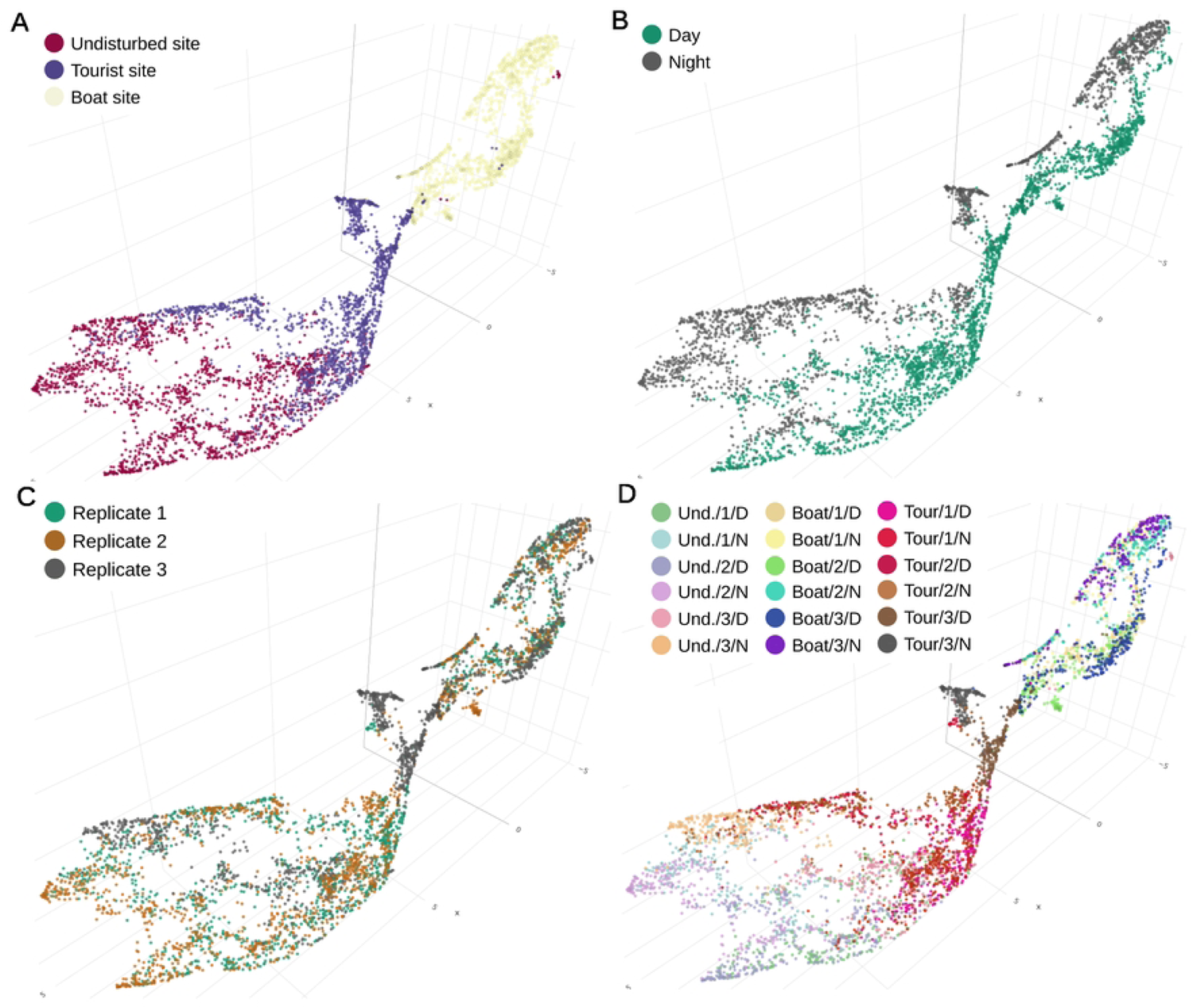
Three-dimensional visualization of the acoustic space of coral reef soundscapes using *CoralSoundExplorer* software. Each dot corresponds to 15 seconds of sound recording. Recordings were made at three sites on the Bora-Bora reef (undisturbed site, tourist site and boat site), over 24-hour periods, defining a day period and a night period, repeated over 3 non-consecutive 24-hour days (replicates 1, 2 and 3). In panels (A), (B) and (C), each sound is colored according to one of these predefined labels. Panel (D) combines the three labels using the following nomenclature site/replicate/period (site: undisturbed as Und., tourist as Tour., and boat as Boat, replicate number 1, 2, or 3, day and night as D or N) to form 18 predefined clusters (3*2*3 = 18). These graphical representations make it easy to explore soundscapes and qualitatively grasp their correspondence with predefined labels. *CoralSoundExplorer*’s interface is interactive, allowing the 3D representation to be oriented as desired and zoomed in on areas of interest. Each dot is associated with the corresponding sound recording, which can be listened to and exported with a click.

In Fig 10A, the color of the dots identifies the recording sites. The three sites appear to have related soundscapes, i.e. they share certain sounds, but they are different, as the overlap between the soundscapes of the sites remains limited. The soundscape of the boat site, for example, shows no overlap with the soundscape of the undisturbed site. The soundscape of the tourist site, on the other hand, features numerous sound points in a region of the acoustic space also occupied by sound points from the undisturbed site. Some dots from the tourist site are also found in the cloud of boat sounds. The distribution of points may be partly explained by the presence of boats: there is a positive gradient from the undisturbed site to the boat site, which explains why the tourist site shares points with these two sites, having periods with and without boats.

Fig 10B shows the difference between daytime and night-time sound environments on the reef. This dichotomy between day and night can be seen at all three recording sites. However, a closer look at the distribution of labeled day and night points for each site reveals subtle variations between sites. A significant region of the acoustic space occupied by the undisturbed site sounds is thus a mixture of day and night labeled points. Sounds common to both periods suggest the activity of animal species during extended periods of dawn or dusk. The distinction between day and night sounds is clearest at the boat site. Here, the daytime soundscape is strongly affected by boat traffic, and contrasts with the nocturnal soundscape, which is certainly much quieter.

In Fig 10C, the dots representing the sounds are colored according to replicate. There is no obvious structuring in the acoustic space, particularly for the boat site and the undisturbed site. This suggests that the sound environments of these two sites remained relatively homogeneous between the recording sessions (which were not consecutive - see Methods). However, this homogeneity between replicates is not found for the tourist site, where the third replicate seems to stand out from the first two.

In Fig 10D, the dots are colored according to a composite label created from the three site/period/replicate labels. This makes it possible to assign identities to the sounds samples according to the 18 modalities of this label, confirming in particular the singularity of the third replicate of the tourist site (clusters Tour./3/day mixed with Boat/all replicates/day).

### 2. Quantification of acoustic similarity between recordings using Silhouette Indices

To quantify acoustic similarities and dissimilarities between groups of sound samples defined by the modalities of a categorical label (Fig 10), *CoralSoundExplorer* proposes the calculation of silhouette indices (Fig 11). For the analysis proposed here, we have chosen to work based on silhouette indices obtained between sites (Fig 11A & S1 Table) and between a composite label made from the 153 combinations of the 18 categories of the composite label (site, day/night, replicate - Fig 11B & S2 Table). Other matrices can be constructed, for example by analyzing silhouette indices between replicates or day/night periods only. A silhouette index tending towards 0 indicates a high degree of acoustic similarity between recordings, reflecting a high degree of overlap of recording clouds in acoustic space. When it tends towards 1, it means that the two categories of recordings do not share common acoustic characteristics, i.e., the clouds do not overlap in acoustic space. The data matrices shown in Fig 11 can be exported as .csv spreadsheets.

**Fig 11.**
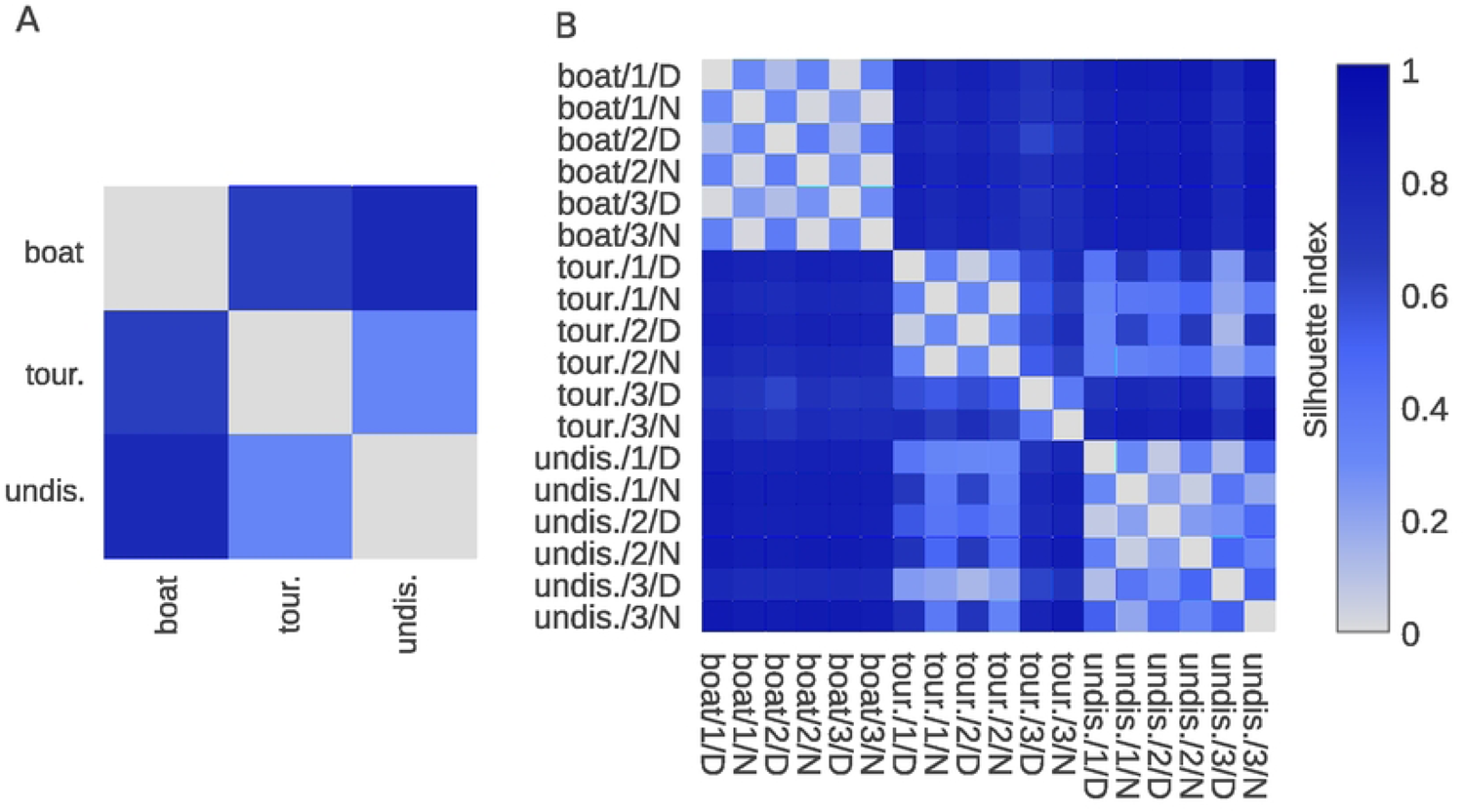
Quantification of acoustic similarity (Silhouette indices) between reef sounds recorded at Bora-bora. Silhouette indices are calculated from 100 UMAPs. The continuous color scale represents the index value (0: the two groups are similar, signifying homogeneous soundscapes; 1: the two groups are completely dissimilar). (A) Recordings are labeled by site only (tour.: tourist, boat: boat site, und.: undisturbed). (B) Recordings are labeled by site, day/night period (D: day, N: night) and replicate number (3 replicates, corresponding to 3 non-consecutive 24-hour recording periods).

Fig 11A shows that the overall differences between sites are not identical. The silhouette indices between the boat site and the others are of the order of 0.7, while the silhouette index between the other two sites is 0.33 (silhouette indices for the comparison between tourist/undisturbed sites: 0.33, between tourist/boat sites: 0.65 and between undisturbed/boat sites: 0.78). This indicates that the soundscapes of the boat site are more different with the other two sites than are the soundscapes of the tourist and undisturbed sites with each other.

By looking at a finer scale of sound groups, following the composite label (Fig 11B), we observe that the silhouette index between day and night on one site are about 0.35 whichever site is concerned (silhouette indices between day and night averaged over the three replicates: 0.35 ± 0.03 for the tourist site; 0.35 ± 0.12 for the undisturbed site; 0.32 ± 0.04 for the boat site; mean ± standard deviation). This value is about the same as the silhouette index for the comparison between tourist/undisturbed sites from Fig 11A but is twice lower than the silhouette indices between the boat site and the two other sites. In other words, it seems that the boat site soundscapes are very different from the soundscapes of the two other sites and that these differences are of a higher degree than the differences between day and night.

In Fig 11B, we also observe low silhouette indices (close to zero) between the soundscapes of the replicates at the boat site, for both day and night. This means that the soundscapes of this site remain similar over several days for both day and night separately (silhouette index for day = 0.08 ± 0.05, for night = 0.02 ± 0.003). This result is slightly less pronounced for the undisturbed site (silhouette index between days = 0.19 ± 0.11, between nights = 0.15 ± 0.09), but the conclusions drawn for the boat site are transposable to the undisturbed site, namely the similarity between nights on the three replicates and between days on the three replicates. For the tourist site, silhouette indices are also close to 0 for replicates 1 and 2, but higher for replicate 3 (Silhouette indices between recordings made during replicate 1 and those made during replicate 2: 0.05 for days, 0.04 for nights. Silhouette indexes between recordings made during repetition 3 and the other two repetitions: 0.59 ± 0.01 for days and 0.63 ± 0.01 for nights). Thus, within a given site, except for the tourist site for the third replicate, the silhouette index matrix quantifies that overall soundscapes are more similar between the three nights or between the three days, than between the nights and days of each replicate. The soundscapes of replicate 3 of the tourist site are about as different from those of the other two sites as they are from those of the same site during the other two recording days.

### 3. Unsupervised identification of acoustic clusters

*CoralSoundExplorer* proposes the use of automatic clustering algorithms to identify groups of recordings sharing common characteristics, without any *a priori* information. This approach is useful to identify clusters of sounds that may be markers of particular sound sources (e.g., specific animals, anthropogenic noises, etc.). For each identified unsupervised cluster, its composition as regards to each predefined label can also be calculated and eventual links between these unsupervised clusters and the period, season or time of the day can be noticed.

Applied to the Bora-Bora recordings, automatic clustering using HDBSCAN with the Excess of Mass (EOM) method identifies two clusters of recordings (Fig 12A). Cluster 0 is constituted at 98.40 % by sounds from the boat site (Fig 12B & S3 Table). Cluster 1 is formed by sounds from the tourist site at 48.87 % and by sounds from the undisturbed site at 49.72 %, making a total of 99.59 %. 16 % of boat site recordings have not been assigned to a cluster (id: −1). This unsupervised clustering indicates that the boat site is acoustically very distinct from the other two sites, confirming the supervised visualization and calculation of silhouette indices.

**Fig 12:**
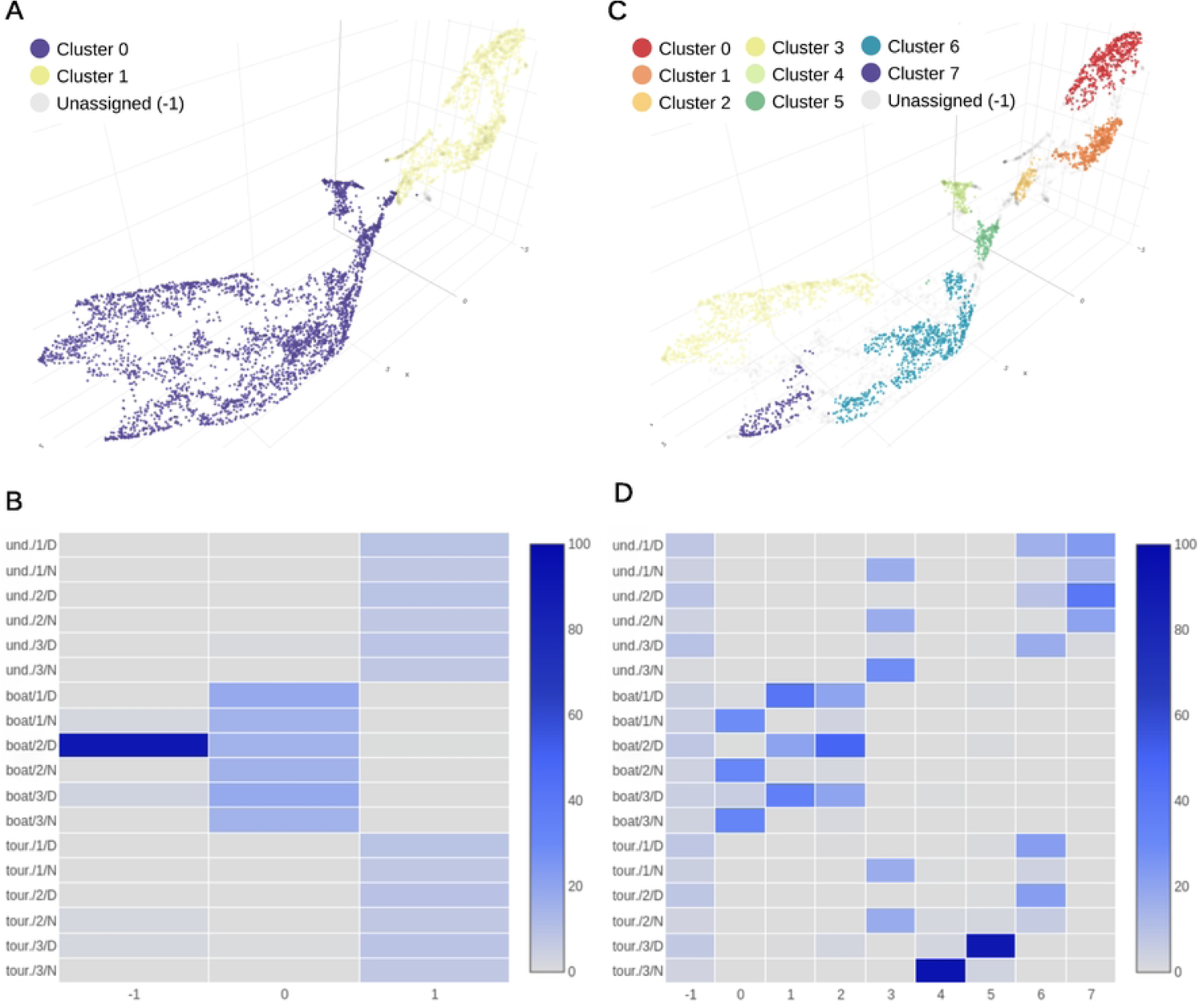
Unsupervised clustering of soundscapes recorded on the coral reefs of Bora- bora. (A) UMAP visualization using HDBSCAN with the Excess of Mass (EOM) clustering method, which identifies two clusters separating the boat recording site from the other two sites (tourist and undisturbed). (B) Contingency matrix of the different modalities of the composite label for each of the two unsupervised acoustic clusters obtained with the excess of mass method (EOM). (C) UMAP visualization using HDBSCAN with the Leaf clustering method. This method identifies eight clusters. These clusters can then be linked to specific sound sources (e.g. boat noise) by manually exploring the recordings (by clicking on the dots, inspecting the spectrograms and listening to the sounds). (D) Contingency matrix of the different modalities of the composite label for each of the eight unsupervised acoustic clusters obtained with the Leaf clustering method.

Automatic clustering using the HDBSCAN with Leaf method identifies eight clusters (Fig 12C). Clusters 0, 1 and 2 shared only recordings from the boat site, cluster 0 was mainly composed of night recordings (93.68 %), and clusters 1 and 2 shared day recordings (respectively 97.86 % and 90.44 % - Fig 12D & S4 Table). After an audio monitoring of these sounds with the *CoralSoundExplorer* audio server, we noticed that these two clusters differ in the nature of the soundscape. Sounds signals contained in cluster 2 show a greater emergence of motorboat sounds than those contained in cluster 1. Tourist and undisturbed sites sounds are both part of cluster 3 (64.95 % composed of undisturbed site recordings and 35.04 % of tourist site recordings) and cluster 6 (44.58 % composed of undisturbed site recordings and 55.41 % of tourist site recordings) that were made up of recordings common to coral reefs. Clusters 4 and 5 were only made up of replicate 3 of the tourist site, respectively night and day time at 94.68 % and 90.54 %. An equal number of recordings from the three sites were not assigned to a specific cluster (unassigned (−1), 36.06 % from the undisturbed site, 29.96 % from the boat site, and 33.97 % from the tourist site).

### 4. Temporal dynamics of Bora-Bora soundscapes and identification of environmental disturbances

*CoralSoundExplorer* offers the opportunity to visualize and quantify how a soundscape varies over time. The temporal trajectories of soundscapes for the undisturbed, tourist and boat sites respectively, for all replicates are plotted over the eight clusters found using Leaf method in Fig 13, 14, and 15. With these displays, it is easy to appreciate the variations in soundscapes over the course of a day, and to see whether these variations are homogeneous between recording days. It is also straightforward to identify events that have disrupted a daily cycle.

**Fig 13.**
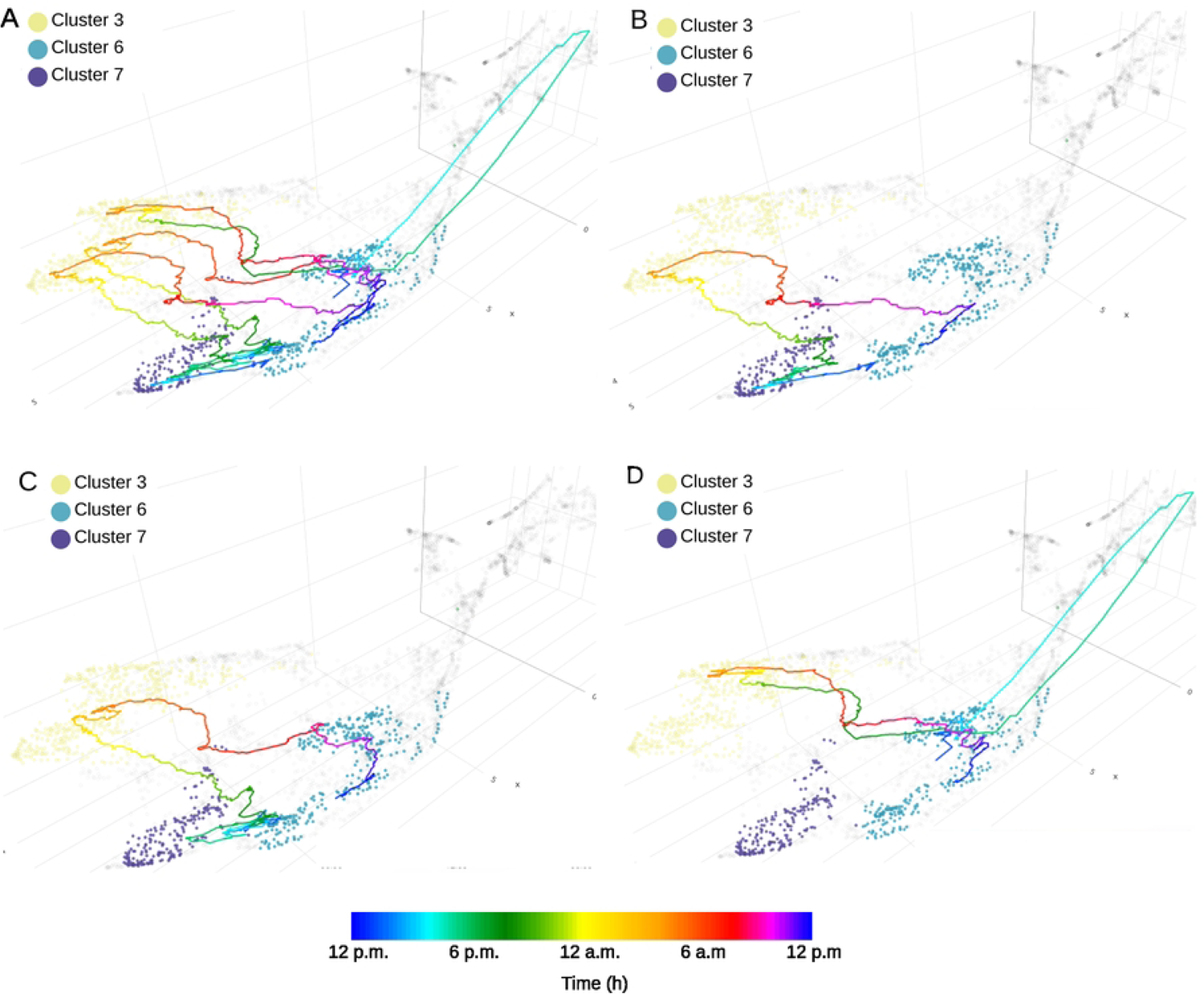
Time path of soundscapes recorded at the Bora-bora undisturbed site. The color scale of the trajectories is related to time. (A) shows the 24-hour trajectory for three different recording days (the three replicates). (B), (C), and (D) show the trajectory for each replicate. Acoustic clusters were generated unsupervised using the Leaf method.

**Fig 14.**
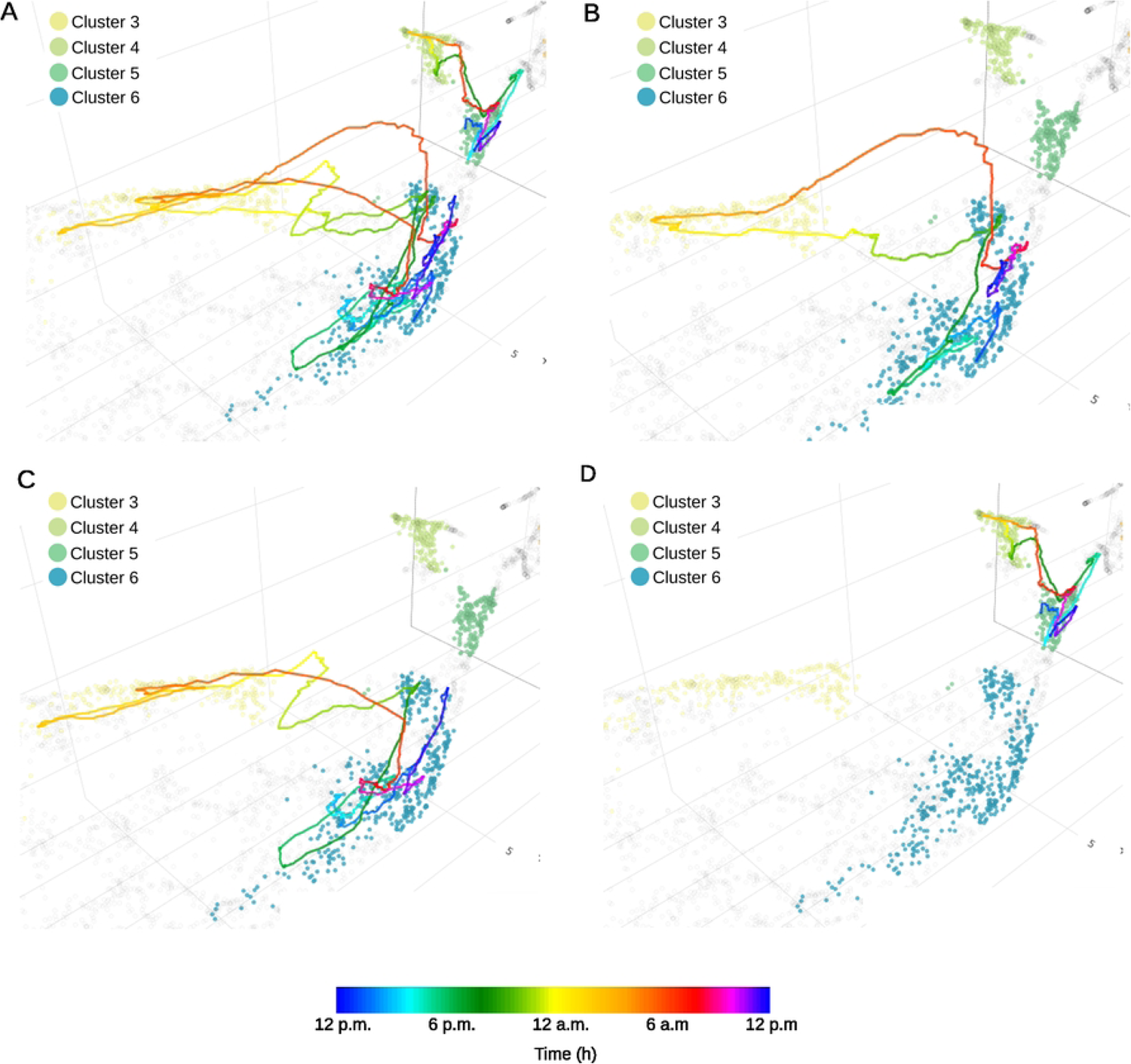
Time path of soundscapes recorded at the Bora-bora tourist site. The color scale of the trajectories is related to time. (A) shows the 24-hour trajectory for three different recording days (the three replicates). (B), (C), and (D) show the trajectory for each replicate. Acoustic clusters were generated unsupervised using the Leaf method.

**Fig 15.**
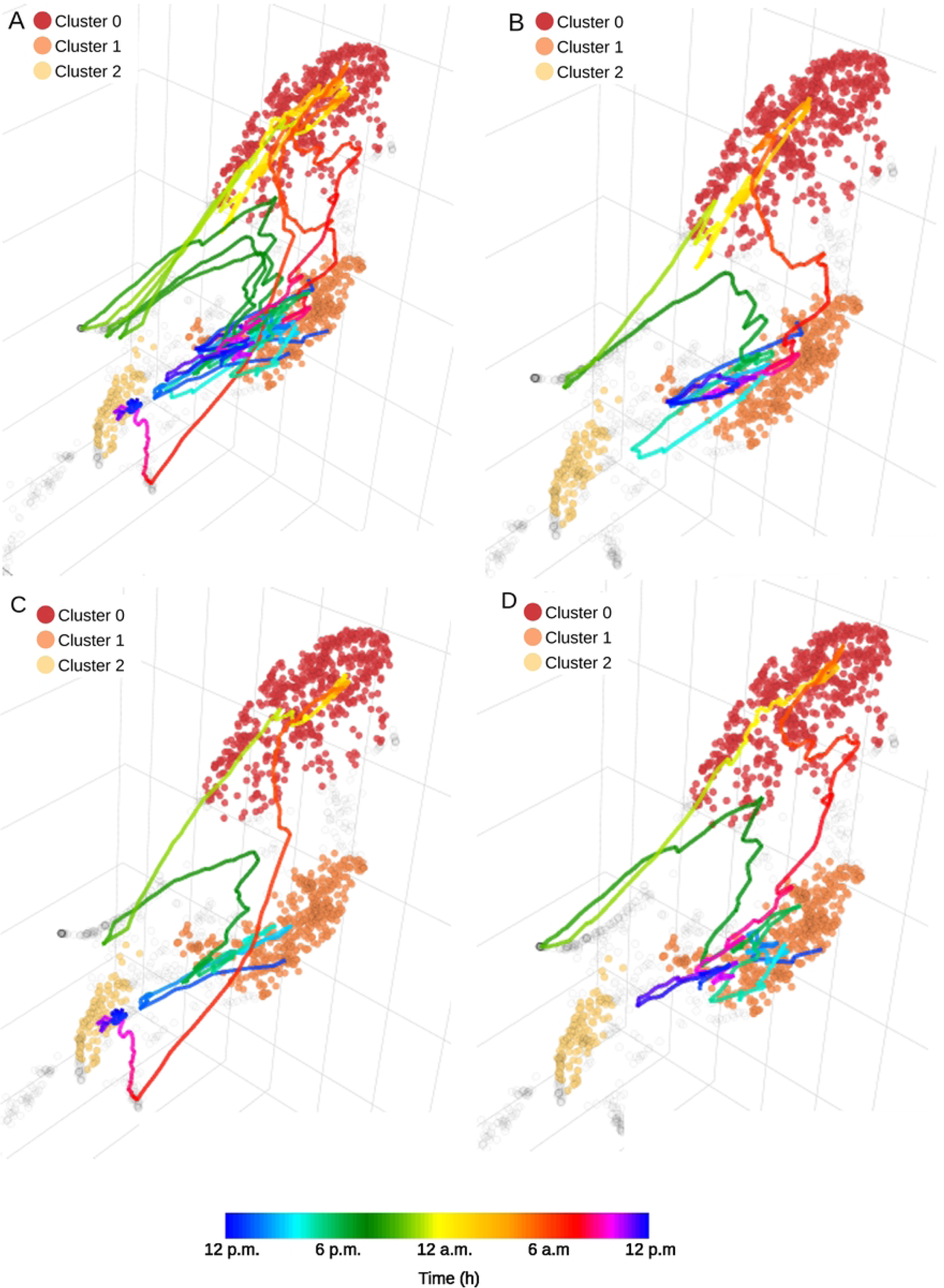
Time path of soundscapes recorded at the Bora-bora boat site. The color scale of the trajectories is related to time. (A) shows the 24-hour trajectory for three different recording days (the three replicates). (B), (C), and (D) show the trajectory for each replicate. Acoustic clusters were generated unsupervised using the Leaf method.

As shown in Fig 13, circadian changes in the soundscape of the undisturbed site are very similar across the three recording days. Clusters 6 and 7 share the daytime recordings of the undisturbed site. More precisely, cluster 6 corresponds to dawn and the beginning of the day, while cluster 7 corresponds to the end of the day and dusk (Fig 13A). Replicates 1 and 2 appear to follow a similar path throughout the day (Fig 13B & 13C). On the third day of recording (replicate 3), the soundscape drifted significantly compared to the first two replicates (Fig 13D). This drift was due to a rain episode lasting about an hour (around 3 p.m.). This episode disrupted the temporal pattern of the soundscape for several hours. As a result, none of the recordings from replicate 3 are to be found in cluster 7, which corresponds to the end of the day and dusk of replicates 1 and 2. The soundscape of replicate 3 returned to a similar pattern similar to the other two replicates during the night and the following morning.

Fig 14 illustrates the temporal trajectories of the soundscape at the tourist site. The soundscape at this site is less homogeneous than at the undisturbed site (Fig 14A). Cluster 3 (representative of nocturnal recordings) and cluster 6 (daytime recordings) are shared with replicates 1 and 2 (Fig 14B & 14C). Other sounds in replicate 3 formed clusters of their own (cluster 4 for night recording and cluster 5 for day recording; Fig 14D).

Fig 15 shows the temporal trajectory of the soundscape at the boat site. During the day, the overall pattern is more difficult to discern than for the other two sites, probably because boat activity is rather chaotic (variation in the number and type of boats, as well as the time of day they sail; Fig 15A). Clusters 1 and 2 (Fig 15B, 15C, & 15D) are common at all replicates but are few related to specific daytime hours. During the late-night hours (from 11 p.m. to approximately 4 a.m.), which correspond to a low boating activity, the temporal trajectories of the 3 replicates all crossed cluster 0. However, the trajectory of replicate 2 deviates from the other two between 10 a.m. and 12 a.m. (Fig 15C), probably highlighting a difference in noise due to marine traffic.

To capture the trajectory of soundscapes in the acoustic space, *CoralSoundExplorer* proposes an original representation showing variations over time in the acoustic distance between the sound and the starting point (time 12 p.m.) of the soundscape’s trajectory. This representation is illustrated in Fig 16. Quantitative data (measurements of acoustic distance versus time) can be exported as .csv files, e.g., for statistical analysis.

**Fig 16:**
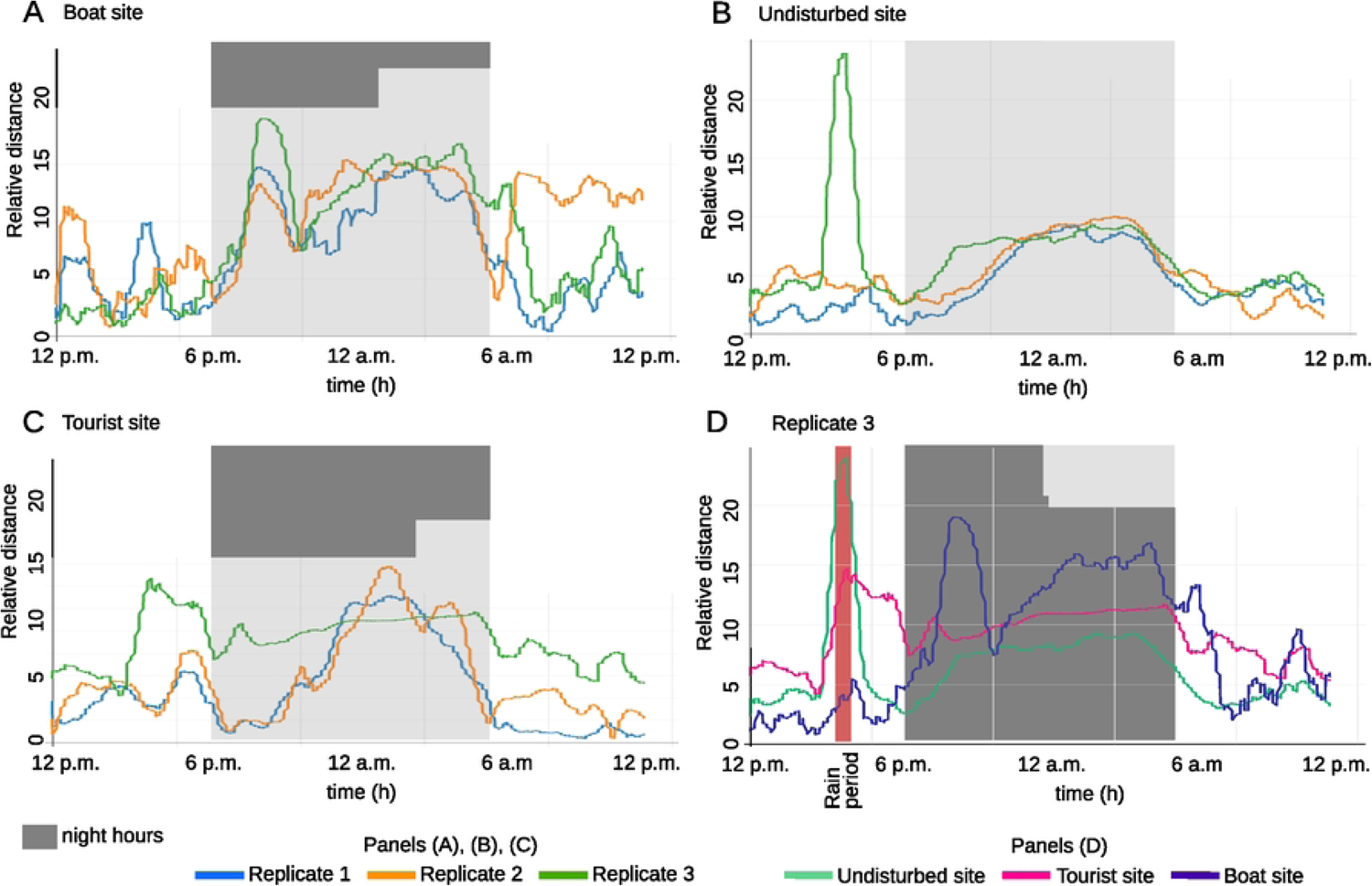
Time trajectories of the soundscapes at the three recording sites. (A) Relative trajectories of the three replicates of the boat site over time. (B) Relative trajectories of the three replicates of the undisturbed site over time. (C) Relative trajectories of the three replicates of the tourist site over time. (D) Relative trajectories of replicate 3 of the three sites over time.

This representation makes it possible to visualize and quantify the temporal dynamics of soundscape variations in an elegant way and using a more stable representation (i.e., based on the 100 UMAPs computations). It is easy to see in Fig 16A that the soundscape of the boat site deviates sharply from its initial position at around 7 p.m., probably due to the sharp drop in boat traffic as night falls. The distance to the initial position remains high throughout the night, suggesting a different soundscape during these hours, then decreases around 8 a.m., but only for replicates 1 and 3.

Fig 16B illustrates the displacement of the soundscape in the acoustic space of the undisturbed site. Again, the distance from the initial point increases at dusk and decreases at dawn. The first two replicates are remarkably similar. For the third replicate, there is a sudden increase in distance at around 3 p.m., indicating that the soundscape changes very rapidly at this time. This disturbance, due to a rainy episode, continues until 3.30 p.m. Only then does the soundscape approach that of the other replicates at the same moment. We can also see that the typical night-time soundscape begins earlier than in the other two replicates, at around 7.30 p.m. This may be due to overcast conditions and the accompanying lower underwater luminosity.

As shown in Fig 16C, variations in distance to the initial point at the tourist site are quite similar to those described for the undisturbed site. Replicates 1 and 2 are relatively similar and replicate 3 was disturbed at 3 p.m. due to the rain. However, unlike the undisturbed site, the tourist site did not recover its initial pattern after the rain: the soundscape is therefore further away from its initial position. It is likely that the rainy event interrupted tourist activity, and durably modified the acoustic pattern.

To facilitate comparison between sites, Fig 16D shows the variations in acoustic distances between soundscapes at the three sites on the third day of recording. Rain had an effect on tourist and undisturbed sites, and a moderate effect on boat site soundscapes, with the main peak occurring at the same time. This effect lasts a few hours for the tourist site while this is not the case for the two other sites. The undisturbed and touristic sites had a greater relative distance during this rain perturbation than the boat site, the latter being deeper, the perturbation and sound masking effect of the rain was potentially less present and also more covered by the noise of the boats.

### 5. Conclusion on the Bora-Bora case study

As illustrated by the study of our Bora-bora database, *CoralSoundExplorer* is a powerful and efficient tool for exploring coral reef soundscapes. This Bora-Bora database had already been explored using conventional methods, such as manual exploration of spectrograms. This exploration, the results of which have recently been detailed [31], had required a considerable amount of time, estimated at around 15 days of work. With *CoralSoundExplorer*, we identified the same features and phenomena in just a few hours, including analysis preparation, computation time and results exploration. *CoralSoundExplorer* therefore makes it possible to rapidly track the dynamics of reef soundscapes over long periods of time.

Saving time is not the only advantage of *CoralSoundExplorer*: the visualization and quantification tools offered by our software make it possible to cluster phenomena that are difficult to detect by manual analysis. For example, the cluster 6 (Fig 12C) corresponds to low-frequency sounds, probably produced by a fish, which is likely a marker for the presence of healthy corals at the tourist site. The cluster 0, again in Fig 12C, also corresponds to sounds emitted by a fish that may only be present at the boat site.

Dynamic variations in soundscapes over a 24-hour cycle are also highly informative. They reflect both the stability of an environment (as in the case of the undisturbed site) and the arrival of particular events (rain), or variations in the soundscape between days (as in the case of the boat site). Subtle changes in the soundscape are thus detected, suggesting the possibility of precise monitoring of the temporal dynamics of the activity of living organisms or other sound sources. The temporal dynamics of soundscapes revealed by *CoralSoundExplorer* provides a tool for rapidly detecting and identifying disturbance and the time required to return to the initial state. For example, rainfall has a lasting effect on undisturbed and tourist sites, while the site with the highest level of human activity, and therefore noise pollution, appears to be less affected.

## IV. Detailed Methodology

This part of the paper first summarizes the analysis principle of CNN/UMAP embedding and general indications about how the semi/unsupervised learning methods work, which form the core of *CoralSoundExplorer*. The problem of UMAP stochasticity is also presented, and its circumvention by using multiple UMAPs is explained. Next, the computational details of the *CoralSoundExplorer* workflow are given, and the parameters of the multiple UMAPs computation process are justified according to dimension reduction validity metrics. Finally, details of the results analysis process not provided in section II are described.

### 1. Soundscape analysis paradigm

The analyses of sound signals proposed by *CoralSoundExplorer* are based on their projections into a representation space derived from a convolutional neural network (CNN) (Albawi 2017) trained for sound analysis. When using a CNN, it is assumed that the projection space, the CNN embedding, is a metric space, i.e., where the notion of distance between individuals projected into this space has a geometric meaning (for the analysis of sound recordings, the individuals are points in the space where each corresponds to a sound signal of a certain duration, decided by the person conducting the analysis). Two points close in distance correspond to two sound signals sharing common acoustic characteristics, and two points further away correspond to two different sound signals. The result is that close sound signals are grouped together in the same embedding subspace and, conversely, groups of different sound signals are moved further apart. This notion of grouping by similarity in the embedded space is that of clusters in machine learning. The relative organization of individuals (sounds) in the embedded space can be studied using a semi-supervised approach. The aim is to find out whether the annotations previously made on each signal, based on knowledge of their recording conditions or content (such as the time and place of recording) enable signals to be grouped (and therefore classified) according to their acoustic characteristics. The relative organization of sounds in the embedded space can be also studied using an unsupervised approach, looking for the most identifiable unsupervised clusters in the embedding space. Explanations for these unsupervised clusters can be found by identifying the similarity of sounds by searching for features that allow grouping or distancing. This can then be sought directly by careful listening and viewing of the spectrogram, or by cross-referencing with labels previously assigned to the recordings. In addition to these analyses based on a categorical organization of the data set, it is also possible to follow a path in the embedding space as a function of a continuous variable value corresponding to each signal. In the context of analyzing the time characteristic of signals and observing the phenology of soundscapes, this continuous variable can be the day or hour of recording, or any other meaningful time interval.

The number of embedding dimensions directly derived from the neural network can be large, posing a problem of interpretability and calculation of inference metrics due to the curse of dimensionality [70]. Cluster extraction or analysis methods can suffer from this. Specifically, density-based cluster extraction methods, such as HDBSCAN, are known to perform poorly in high dimensions (71). To overcome this difficulty, it is useful to use dimensionality reduction methods. The best-known is PCA, but this linear approach is often unsuitable for complex problems due to the potentially significant loss of information at low dimension. Several nonlinear methods have been proposed to overcome this problem, such as t-SNE methods [72] or, more recently, the UMAP approach. The latter is based on a consistent mathematical approach that justifies the globally and locally conservative nature of the data structure. Thus, UMAP generally preserves the clusters present in the dataset on which it is applied and even tends to increase the clustering effect [73]. However, UMAP is often only used as an illustrative support in 2- or 3-dimensions due to the stochastic nature of its algorithmic implementation. This stochastic character, leading to different representations depending on a random seed, cannot be ignored when using UMAP to extract quantitative measurements, and its effect on projection and on the analyses that can be deduced from it needs to be addressed. One solution, which we have employed here, is to work based on several realizations of UMAP from the data and use the central tendencies to characterize and quantify the organization of sounds in the UMAP projection space.

### 2. Signal pre-processing and VGGish embedding

In a first step, *CoralSoundExplorer* segments each raw sound recording into one-second segments (the duration of each raw recording for the Bora-bora dataset was 60 seconds). Each one-second sound segment is then transformed into a log-melogram. The number of Mel bands is set to 64, from the minimum frequency analysis to the maximum one. These limits can be set by the user, for the present study on the Bora-Bora coral reef they were set to 70 Hz and 2 kHz. The number of time steps in the melogram is set to 100, using a 2048 samples STFT Hanning window and a hop size step of 441 samples (Fs /100). The 64×100 values of each melogram amplitudes are transformed using the natural logarithm.

Each resulting log-melogram is then presented as input to the pretrained CNN: the VGGish network. The output of this neural network produces 128 features that are taken as projection vectors for the embedding [51,74]. Each one-second segment of sound is thus represented by 128 dimensions that position it in a kind of acoustic hyperspace. *CoralSoundExplorer* allows to work with integration times longer than one second (which is the case here with the reef recordings, where we have chosen an integration time of 15 seconds). The projections of a defined number of successive one-second sounds (individuals) can be grouped. This is done by calculating the centroids of their coordinates in the embedding space. In the case of the Bora-Bora dataset used as an example in this paper, the original dataset of recordings was composed of 1305 separated recordings of 1 minute, leading to 1305*60=78300 one-second frame individuals in the original VGGish embedding. By grouping the successive one-second frames by 15, the final data set in the final VGGish embedding was reduced to 5220 individuals (each individual being 15 seconds of recording). In addition, we retained only 119 of the original 128 dimensions, as nine dimensions were non-significant (i.e., with constant values over the whole dataset). For each of these 119 dimensions, the embedded dataset was robust scaled, i.e., centered on the median and scaled to the interquartile range between the first and third quartiles, in order to balance the effect of each dimension before UMAP dimension reductions.

### 3. UMAPs computation and choice of UMAPs parameters

From the VGGish embedding dataset, *CoralSoundExplorer* calculates 100 independent UMAP transformations (Python package UMAP-Learn version 0.5.3, https://umap-learn.readthedocs.io). Each UMAP algorithm processing differs from the initialization random seed. The metric used to compute the distance is the Manhattan distance. The number of neighbors is set to 15 and the minimum distance in the UMAP representation is set to 0. The decision to set the final number of UMAP dimensions to 3 and the number of UMAPs to 100 was based on preliminary studies conducted as part of the present work (see below).

#### 1. Number of independent UMAP computations

We have determined the number of independent UMAP processes required to smooth the stochastic effect of UMAP projections using averaged distance matrices. The choice of using distance matrices to assess the stability of metrics derived from UMAP stems from the fact that the absolute values of the coordinates of UMAP projections make little sense for studying the relative organization of individuals (i.e., sound samples). Only relative distances are necessary. Indeed, most of the analysis proposed here, except for the ‘path’ visualization, are based on a distance matrix and not on coordinates.

For each UMAP realization, the distance matrix of the individuals was computed using the Euclidean distance, since the UMAP space projection is Euclidean. For the *n*^th^ UMAP projection, an average distance matrix was then computed from the distance matrix of the *n*^th^ UMAP projection, and the previous *n*-1 distance matrices. To assess the stability of the *n*^th^ average distance matrix as a function of *n*, we calculated the relative mean absolute difference and the relative maximum absolute difference between the pairwise values of the *n*^th^ average distance matrix and the *n*-1^th^ average distance matrix. These difference metrics are related to the mean distance values of the *n*^th^ distance matrix values. Due to the symmetric property of the distance matrices and the zero values in their diagonals, only the lower (or upper) triangle of these matrices without their diagonals was used. The formula for these metrics is as follows:

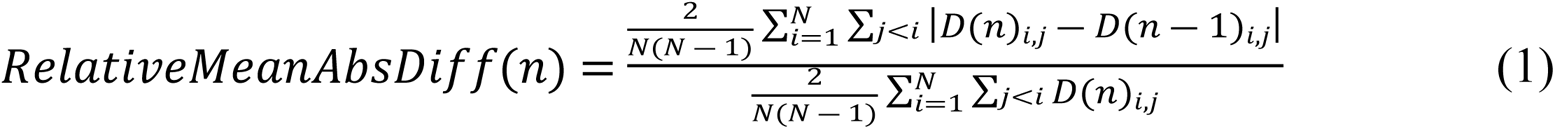

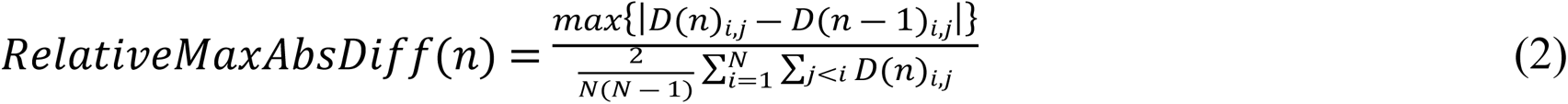

where D(n)_i,j_ are the values at row *i* and column *j* of the average distance matrix made from *n* individual matrix distances, and *N* is the total number of rows or columns of the distance matrix. The number of realizations (equal to the number of distance matrices used for averaging) varied from 1 to 100, and the final number of dimensions for the UMAP computation was either 119, 5, 3 or 2. The dimensionality of 119 is the same as that of the initial VGGish embedding, considering only those dimensions that actually support the information, so that in this case the UMAP algorithm does not strictly perform a reduction but simply transforms the space according to the UMAP principles. The dimensionalities of 3 and 2 allow us to visualize the embedding. The dimensionality of 5 is an intermediate representation intended to be sufficient to suppress the “curse of dimensionality” effect while keeping enough information.

The results are shown in Fig 17. Whatever the number of UMAP dimensions used to calculate the distance matrices, the relative mean and max absolute differences between the average matrix *n* and *n*-1 decrease exponentially with *n,* demonstrating the convergence of the average distance matrix as the number of multiple UMAP transformations increases. This result is an example of the application of the central limit theorem which, assuming that the positions of individuals in UMAP projections are bounded and therefore also the values of the distance matrix, states that this convergence should follow a 1/√n law. The issue here was therefore more to assess convergence speeds and their stabilities than to establish the convergences themselves. For each UMAP dimensionality, the maximum relative difference between the average distance matrices obtained from 100 and 99 distance matrices is less than 0.05%, whereas the same metric obtained from the first and second distance matrices was greater than 1%. The convergence speeds of this metric as a function of *n* appear to be roughly the same, with perhaps a very small advantage for the cases of higher dimensional UMAPs. The stability of the convergences of the UMAPs mean distance matrices with *n* does not seem to be equal regarding the UMAPs dimensionality, in particular the 2d UMAPs which provided a less smooth decrease in the relative mean error curve. If we consider the maximum relative error curves, the decay rates also appear to be the same whatever the UMAPs dimensionality, and each of these curves leads to a final value of less than 1%.

**Fig 17:**
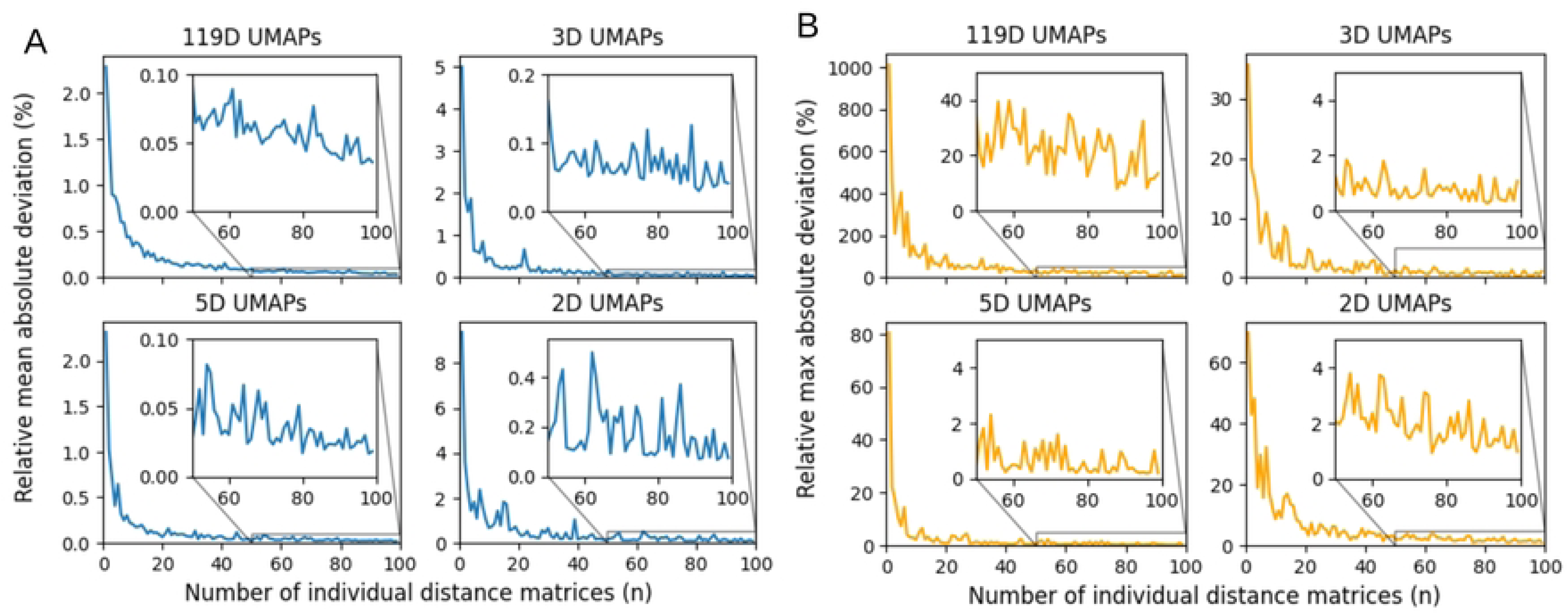
Absolute differences between distance matrices computed from n-1 UMAPs and n UMAPs. Absolute differences between distance matrices computed from n-1 UMAPs and n UMAPs relative to the mean distance observed in the distance matrix computed from n UMAPs. (A) Mean absolute differences. (B) Maximum absolute differences.

#### 2. Number of UMAPs dimensions

A systematic question that needs to be asked in the case of dimension reduction, using UMAPs or any other dimension reduction technique, concerns this final number of dimensions: how many dimensions are sufficient to retain most of the information? To answer this question, we have calculated two metrics of topological conservation properties as a function of UMAP dimensionality. The first, more related to the local conservation property, is the Degree of Local Preservation (DLP) [75]. The second, related to the global conservation property, is Random Triplet Accuracy (RTA) [76]. For these two metrics, the following values were obtained from average distance matrices processed from 100 independent UMAPs computations.

The DLP metric measures the average rate of nearest neighbors retained after dimension reduction on a number of *n* points. This metric depends on the number of neighbors considered *k* and is obtained from equation (3). Where *knn_UMAP i_* are the nearest neighbors of point *i* in the UMAP projection and *knn_VGGish i_* are the nearest neighbors of the same point in VGGish space.

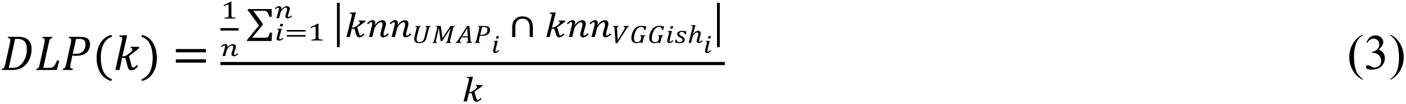

Given the small size of the coral reef dataset used here, this metric could be calculated using the total number of individuals *N* as *n*, which would avoid the stochastic aspect due to the choice of individuals *i*. This metric must be evaluated in relation to its ‘by chance’ value, which depends on the number *k* relating to the number of individuals in the entire data set. This ‘by chance’ value is given by equation (4). We can also use a Monte-Carlos simulation, especially when *N* is large, as binomial coefficients are difficult to compute in such cases. This Monte-Carlos simulation consists of random permutations of individual positions in the original or final projection space before the distance matrix computation, and then calculating the DLP(k).

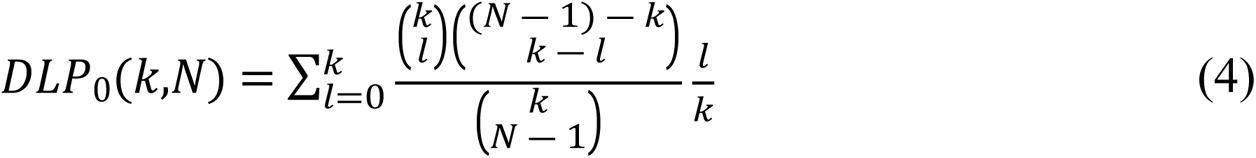

Fig 18 shows the DLP and the DLP rectified from its ‘by chance’ value as a function of the number of nearest neighbors considered for the 4 UMAPs dimensionality values (119,5,3,2). DLP is strongly influenced by the number of nearest neighbors considered, but the effect of the dimensionality is marginal. The maximum values of the rectified DLP curves and their approximate averages are 0.66 and 0.39 for 119d, 5d and 3d UMAPs and 0.65 and 0.39 for 2D UMAPs.

**Fig 18:**
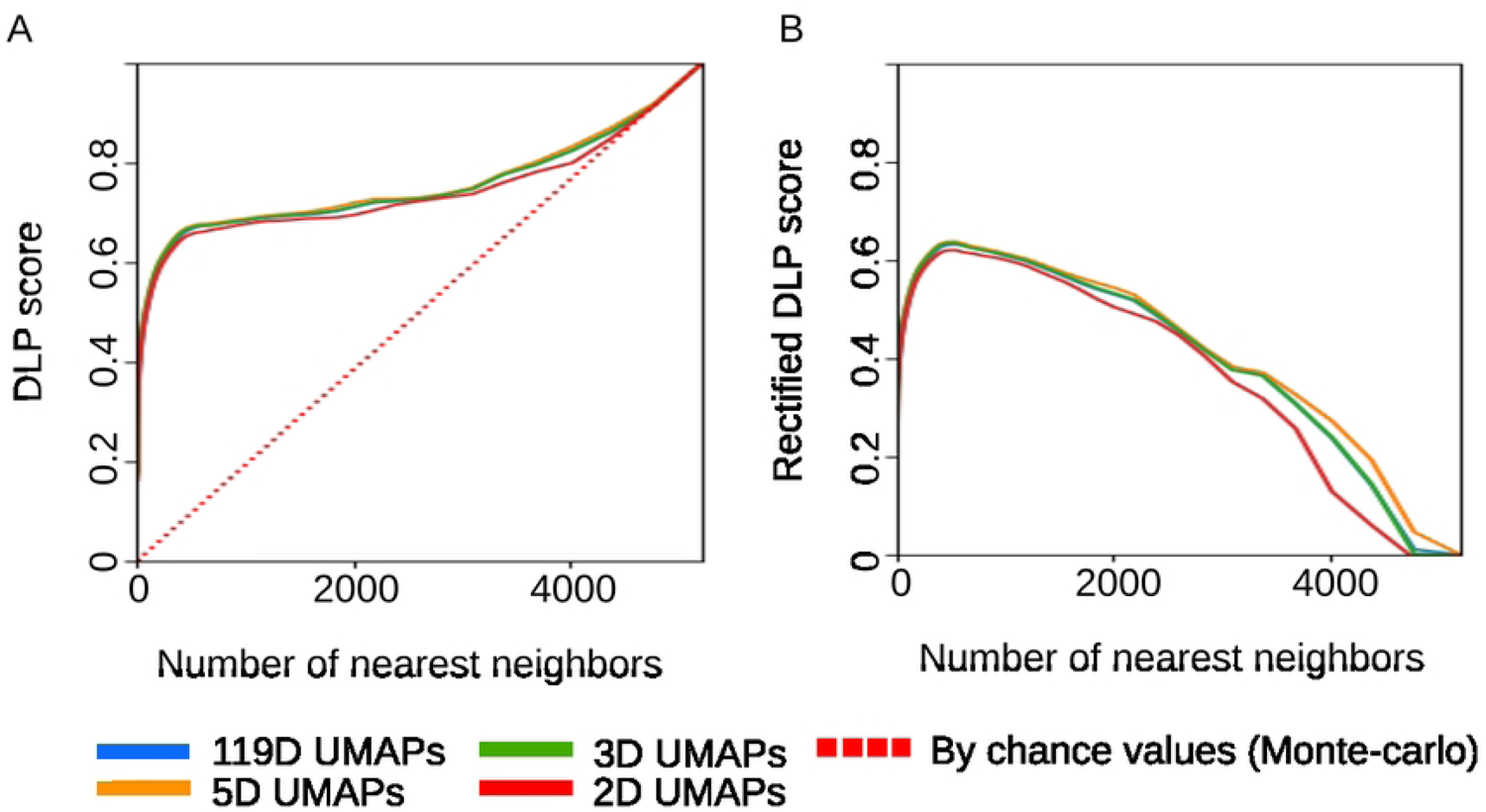
Local topography preservation metric of the mean average distance matrices computed from UMAPs as a function of the number of nearest neighbors for all projected individuals. (A) Degree of Local Preservation (DLP) and (B) its rectified value by its ‘by chance’ value.

The RTA metric considers random triplets of individuals and measures the ratio of these triplets whose inter-individual distances ranking are the same in the original space and in the projection space. This metric is stochastic, and a large number of triplets must be used to make it converge. The ‘by chance’ value of the RTA is equal to (3*!)^-1^*.

Fig 19 shows the RTA scores as a function of UMAP dimensionality and its ‘by chance’ value. The number of triplets used for each average distance matrix was set at 1000, and 1000 RTA processes were performed to compute the boxplots in this figure. A one-way ANOVA performed on this dataset indicated a significant difference related to UMAP dimensionality: *F*(3.996) = 15.8; *p*<0.001. Tukey’s HSD pairwise group comparisons tests yielded p-values greater than 0.01 only between 119D UMAPs and 5D UMAPs and between 3D UMAPs and 2D UMAPs. These tests also indicated that the two UMAPs with the highest dimension performed better in the sense of RTA. However, this effect was small, less than 0.01% on the RTA.

**Fig 19:**
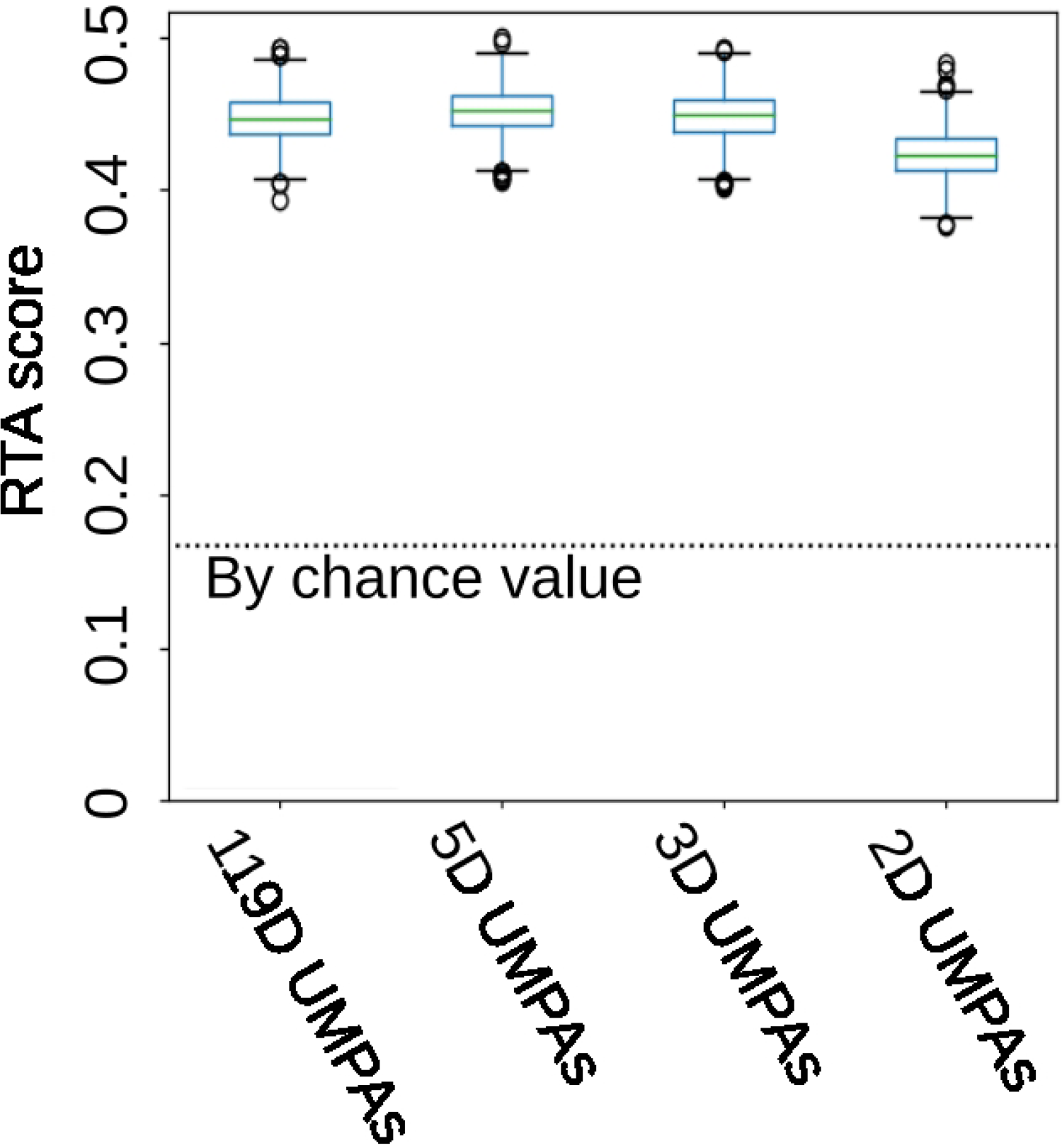
Global topography preservation metric of the mean average distance matrices: Random Triplets Accuracy (RTA) and its ‘by chance’ value computed from UMAPs. The boxplots were drawn from 1000 independent RTA computations for each UMAPs dimensionality. Each individual RTA value was computed from 1000 triplets.

The pilot study on the number of UMAPs required to make the distance matrix converge showed that from 100 individual UMAP realizations, the average error becomes close to zero, and that this metric is the most stable for the 3 largest dimensionalities considered (119D, 5D and 3D). The study on the information retained after UMAPs transformation showed no obvious differences depending on the dimensionality considered. This result of relative independence of UMAP dimensionality on final topography is like those of a previous study in the context of using UMAPs to study communications in cetaceans [77]. Thus, any of the 3 highest dimensionalities considered could be used to compute the 100 UMAPs. As 3D UMAP transforms are less resource-intensive and can be visualized, this was the choice made.

#### 3. Dataset Analysis process

As previously mentioned, we propose the silhouette index to test whether predefined labels can be used to discriminate individuals. The silhouette index is directly linked to the notion of topography, which lies at the very heart of clustering and UMAP transformation. The computation of the silhouette index has been shown to be compatible with UMAP transformation [78]. Conceptually, for each individual, this index is related to the ratio between the average distance separating it from other individuals belonging to the same category and the average distance separating it from individuals belonging to the closest category. For a category, the silhouette index is the average of all the individuals in the cluster. Silhouette index values could occur, which theoretically means that sounds are assigned to the wrong modality. In the case of the present use of the silhouette index, this does not really make sense. However negative values, slightly less than zero, could appear. These should be treated as if they were zero. For the computation of the silhouette index proposed here, only two categories are taken into account at a time, and the average distance matrix of the 100 3D UMAPs used is that obtained in the *Calculation of UMAPs and setting of UMAP parameters* section of this document.

To find unsupervised clusters in the dataset, we chose to use the HDBSCAN algorithm [79]. The use of UMAPs as the basis for this cluster-finding process is motivated by the fact that the UMAP process is known to increase the clusterability and therefore the HDBSCAN performance [78]. Indeed, the average Hopkins statistics [80], which assesses the clusterability of a dataset was 0.98 for the 3D UMAP used to calculate the average distance matrix, while it was 0.79 for the 119d VGGish space. The input to the HDBSCAN algorithm is the same average distance matrix used to calculate the silhouette index. The HDBSCAN implementation is that available in the HDBSCAN* Python package (version 0.8.33 https://hdbscan.readthedocs.io). The minimum number of samples to consider a cluster and the minimum number of neighbors to consider an individual as a core point were set to the same value of 100, which, in terms of recording duration, correspond to 25 minutes. The “cluster_selection_epsilon” and “alpha” parameters, which are linked to the merging of individuals into clusters, were left at 0 and 1 respectively, their default values. The algorithm used for final cluster selection is either the “Excess of Mass” algorithm or the ‘leaf’ algorithm. The former is supposed to lead to larger and fewer clusters, while the latter is supposed to give the most fine grained and homogeneous clusters according to the HDBSCAN* Python package presentation (https://hdbscan.readthedocs.io/en/latest/parameter_selection.html accessed Nov. 13th 2023). In this way, two different clustering are obtained at 2 different scales.

*CoralSoundExplorer* allows to visualize temporal paths linking points representing sounds in acoustic spaces defined by 2D or 3D UMAP representations. In the example of the Bora-Bora recordings considered in the present study, it is thus possible to trace these paths over the course of a day. These paths are calculated as follows. The individuals (or the sound projections) corresponding to each day recording session at each recording place are first sorted by date, from the first recording date (noon) to the last recording date (noon the following day). From the sorted individuals, the coordinates of each path point were obtained using a sliding average over a one-hour frame to reduce the noise effect. The central time of the rolling mean frames is taken as the time point. This process is applied to each of the 100 3D UMAPs to obtain a stable and reliable representation of the soundscape phenology. The distances between an average starting point and each of the path points computed on the 3D UMAPs are then measured. For each of the three sites considered, the average starting point is computed from the coordinates of each of the start points on each of the three recording days (at t = 0). This measure is related to the average distance between the starting point and its 100 first nearest neighbors. For each day and each site, *CoralSoundExplorer* retains the median of 100 measurements (one per 3D UMAP) of distances relative to the average starting point as a function of daytime, leading to a single stable measurement.

## V. *CoralSoundExplorer*: Main achievements and limitations

Although there are many tools and metrics available today for analyzing soundscapes, until now there has been no software capable of visualizing them in a 2- or 3-dimensional space while associating relevant quantitative measurements. *CoralSoundExplorer* is the answer to this need. Suitable for non-programming-expert users, while remaining open to modification by experts, *CoralSoundExplorer* is a tool for efficiently and rapidly detecting sound events in reef soundscapes. A powerful visual and quantitative tool, *CoralSoundExplorer* provides a low-cost solution to coral reef monitoring and management problems.

### 1. A powerful software tool for visualizing and quantifying coral reef soundscapes

Based on a neural network directly trained and fed by sound or one of its classical 2D spectrogram transformations, *CoralSoundExplorer* avoids the often time-consuming and ambiguous process of extracting conventional acoustic indicators, as is typically the case in soundscape studies [40]. The use of CNNs trained directly on spectrograms and fed with spectrogram data thus provides better results for the analysis of coral reef soundscapes [81].

*CoralSoundExplorer* is structured into two distinct parts: the resource-intensive computational part, which concerns pre-processing, VGGish and UMAP projections and the calculation of all results, and the visualization part, which uses a lightweight dedicated interface, directly available in a web browser but which does not require an internet connection when in use. The calculation part is fed by sound files and requires the user to fill in a spreadsheet in ODS or XLSX format. *CoralSoundExplorer* provides a template for this spreadsheet, which must list all original files, their predefined labels and computation parameters (default values are proposed). The calculation section produces a data file (HDF5 format, [82]) containing all the results of the computational part and used to display the results. The functional separation between calculation and results visualization enables tasks to be shared between different people working on the same project.

The visualization tool uses color-scaled matrix displays, 2- or 3-D map visualizations of colored points and 2-D metric graphs. The sound signals corresponding to each point on the 2- or 3-D visualization map can be listened to and their spectrograms visualized directly via the *CoralSoundExplorer* interface after connection to the audio module. These features make *CoralSoundExplorer* an immediately usable and easy-to-use tool for the visual analysis of sounds from a field recording campaign without extensive knowledge of machine learning. For instance, visualizing the phenology of the soundscape through paths in a 2- or 3-D representation enables a monitoring operator to easily identify aberrant paths. In addition to visual representations, the results of the various analysis processes (matrix values, unsupervised cluster id, etc.) can be exported to a text file for external processing.

### 2. Technical choices inevitably involve compromises

The choice of using 100 UMAPs to obtain convergent quantitative results is quite strict here in terms of permitted deviation from the mean and maximum errors of mean distance matrices. If the data to be processed are large, UMAP calculations can be time-consuming. If this is penalizing, then it is possible to relax the constraints on the number of UMAPs to be averaged without leading to a significant change in the conclusions of the analysis.

Another limitation is that the analyses performed here on the number of UMAP dimensions and the number of UMAPs are only really valid for the dataset used. Although coral reef soundscapes all share common characteristics, it is possible that for datasets larger and more complex than the CNN VGGish datasets, the values of 3 dimensions and 100 UMAPs are not suitable for a good representation of data organization and convergence of these representations. A higher UMAP dimensionality may then be required, and the UMAP hyperparameters will have to be modified.

UMAP is an effective method of dimensionality reduction, which is why we have chosen to use them for *CoralSoundExplorer*. However, UMAPs are not perfect in terms of topography preservation: here for example, RTA and DLP values are of the order of 40% and 50%. The expert user of *CoralSoundExplorer* will eventually be able to implement other methods based on the same principle but with a different optimization method that could be more efficient, such as triMAP or pacMAC [76].

*CoralSoundExplorer* uses the silhouette index to analyze the validity of using manual labels for data partitioning. Other indices could also be used. Some indices may reflect notions of clusterability different from that of the silhouette index, which is based on a notion of relative proximity. The Density-Based Clustering Validation index [83] is thus based on the notion of density and its variation between clusters. The use of a support vector machine approach [84] coupled with the adjusted mutual information index [85] could further reflect a notion of boundaries between clusters. These other indices are not implemented in *CoralSoundExplorer* but could easily be by a more experienced user.

For the unsupervised cluster search step, one difficulty arises from the fact that the HDBSCAN algorithm can give different results depending on its parametrization. his parametrization cannot therefore be carried out completely blindly. The results given by HDBSCAN must always be evaluated and compared with the reality in the field. The two different modalities of the final clustering algorithm used here give results consistent with the spatio-temporal analysis of the recordings. It is possible, however, that finer or coarser clustering would have made sense with a different parametrization, or, on the contrary, produced meaningless sub-cluster partitions. To be sure of the meaning of a cluster, it will always be preferable to go back to listening to the sounds or visualizing their spectrogram of each cluster.

*CoralSoundExplorer*’s monitoring of sound evolution over the course of a day and in a given location is based primarily on a visual study of the paths in a 2- or 3-D representation of UMAP space. This visualization enables an initially qualitative assessment. In quantitative terms, *CoralSoundExplorer* proposes a metric based on the extremes of the dispersions of the differences between two paths at the same time. Other metrics could be developed or derived, e.g., from the field of GPS trajectory analysis [86] or the field of time series analysis [87].

In summary, *CoralSoundExplorer* provides a novel software foundation, enabling the global exploration of coral reef soundscapes by non-programming-expert users through a user-friendly interface. As *CoralSoundExplorer* is open-source software, written in the Python language, both the calculation and visualization parts can be adapted by the user to specific research needs, even outside the framework of coral reef soundscapes. For example, the VGGish model used here to generate the CNN integration can be replaced by another CNN better suited to the intended task, and new analysis functions can be added. Our research team is thus already developing a version of *CoralSoundExplorer* fitted for terrestrial soundscapes and another one for shorter recordings. Although *CoralSoundExplorer* currently relies on the analysis of archival data, its workflow could be adapted to real-time monitoring of soundscapes, reducing the weight of storage data and providing useful alerts for instant reef management.

## VI. Software installation procedure and instructions

*CoralSoundExplorer* provides an efficient environment for working with acoustic data. Here we give a quick overview of how to install and use the software. Its source code and the dataset used for illustrative purposes in the present paper (Bora-Bora) are available online on GitHub https://github.com/sound-scape-explorer/coral-sound-explorer.

### 1. Supported Platforms

*CoralSoundExplorer* is compatible with various operating systems capable of running Python 3 and Node.js. We have extensively tested the software on the following platforms:

1. **Windows**: Supported on Windows 10 and later versions.
2. **macOS**: Compatible with the latest M1 and M2 chips.
3. **Ubuntu**: Installable on Ubuntu 20.04 and newer.

### 2. System Requirements

Prior to installation, ensure that your computer meets the following requirements:

- A minimum of 4 GB of RAM for optimal performance.
- A multi-core processor (2 cores or more) for efficient audio processing.
- About 15 GB of free disk space for installation of dependencies.
- Sufficient storage space for audio files and processed data.
- For optional accelerated audio processing, we advise using an NVIDIA GPU.

### 3. Installation Instructions

To install *CoralSoundExplorer* on your machine, please refer to our detailed installation guides available here https://sound-scape-explorer.github.io/docs/CSE/installation/.

### 4. Software Architecture

*CoralSoundExplorer* consists of three distinct modules: Campaign, Processing, and Visualization, tailored for campaign creation, settings customization, and data sharing (Fig 20). These domains can be used independently, accommodating various usage scenarios.

**Fig 20:**
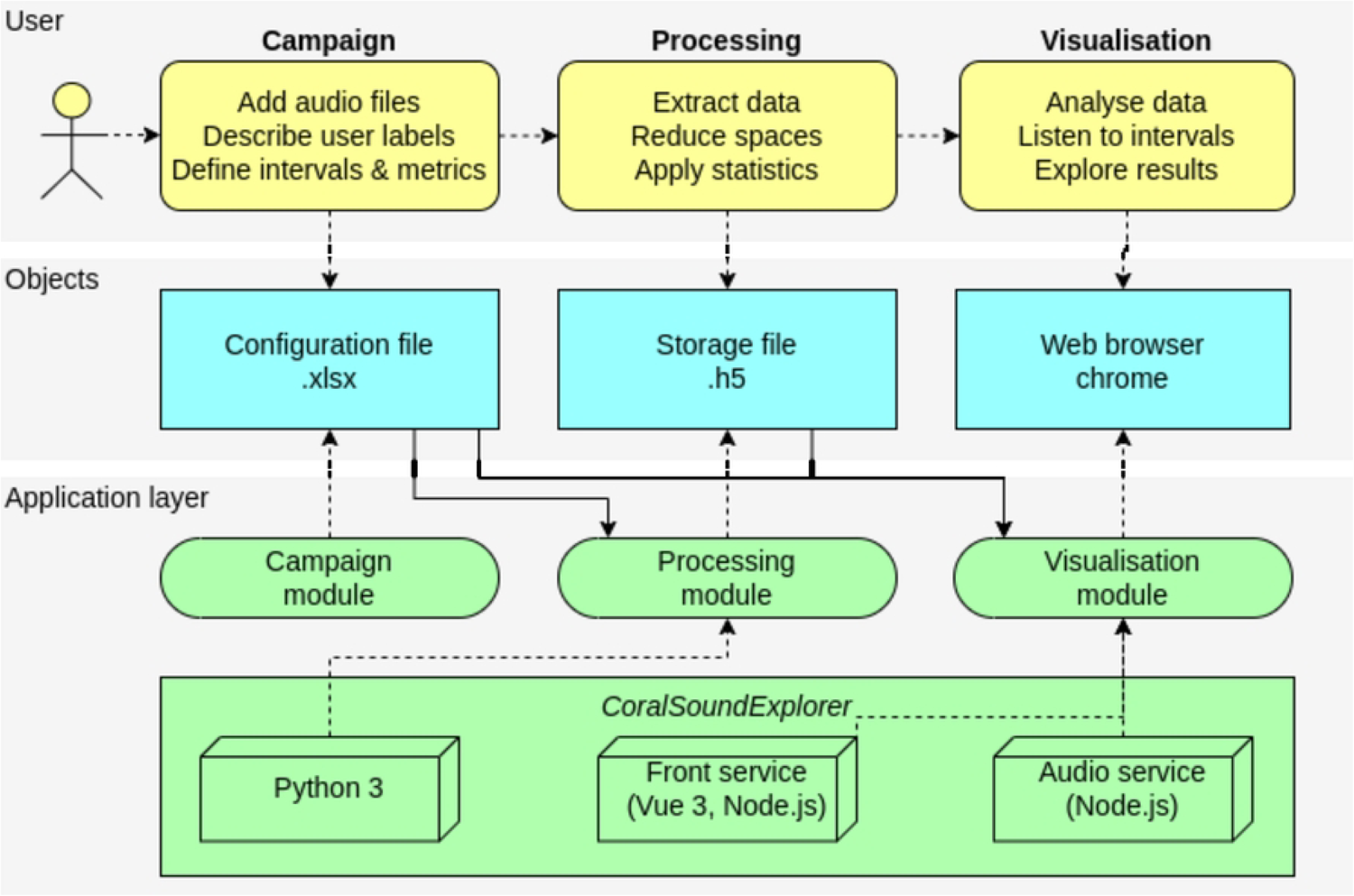
Architecture of *CoralSoundExplorer* software. The top panel describes the user actions. The second panel presents application objects the user interacts with. The bottom panel displays the application modules with underlying technologies.

#### 1. Campaign Module

Upon collecting field audio recordings, researchers can define configuration files and run new campaigns using *CoralSoundExplorer*. Detailed instructions can be found in the campaign documentation https://sound-scape-explorer.github.io/docs/CSE/modules/campaign/.

#### 2. Processing Module

The Processing Module is the core of *CoralSoundExplorer* and is written in Python 3. It uses a series of algorithms to extract valuable metrics from audio recordings. The user must provide the system with a configuration file in .xlsx format, which guides the computation process by indicating the audio file paths and the necessary settings (default parameters are proposed). The generated data is stored in .h5 format, enabling seamless integration with the Visualization Module and facilitating data sharing between users.

The Processing Module is the most important component of *CoralSoundExplorer*, both in terms of size and performance requirements. Processing times for the Bora-Bora dataset taken as an illustrative example in this paper range from 40 minutes (on a GPU-accelerated standard desktop) to 3 hours (on a standard laptop with a CPU). For additional details, please consult the Processing documentation https://sound-scape-explorer.github.io/docs/CSE/modules/processing/.

#### 3. Visualization Module

Implemented in TypeScript using Vue 3, the Visualization Module provides an intuitive and interactive graphical interface with an additional, independent Audio service. This web application operates locally, eliminating the need for an internet connection. A public instance is also available online at https://sound-scape-explorer.github.io/docs/CSE/extras/visualisation-online, enabling basic exploration without any installation. Users can input h5 storage files generated by the Processing Module (including files they have not generated themselves but have shared with other users). The Visualization Module offers a comprehensive set of features for visualizing and listening to processed results, empowering researchers to gain insights and make data-driven decisions.

The Audio service is a lightweight web server designed to seamlessly integrate with the Front service (web application). It enables real-time playback and spectrum analysis of audio files.

For a detailed guide on using the Visualization Module, please refer to our documentation https://sound-scape-explorer.github.io/docs/CSE/modules/visualisation/.

### 5. Future Development

To facilitate bug fixes and the addition of new features, we encourage users and researchers to share their feedback and suggestions by creating new GitHub issues. Please read our contribution guidelines https://sound-scape-explorer.github.io/contributing. Your input is invaluable for the continued improvement of *CoralSoundExplorer*.

## Supporting information

**S1 Table. Silhouette indices between recordings grouped by site only** (tour.: tourist, boat: boat site, und.: undisturbed) (see Fig 11A).

**S2 Table. Silhouette indices between recordings grouped by the composite label site/period/replicate:** site (tour.: tourist, boat: boat site, und.: undisturbed), day/night period (D : day, N : night) and replicate number (3 replicates, corresponding to 3 non-consecutive 24-hour recording periods) (see Fig 11B).

**S3 Table. Contingency values of the different modalities of the composite label for each of the two unsupervised acoustic clusters obtained with the excess of mass method (EOM)** (see Fig 12B).

**S4 Table. Contingency matrix of the different modalities of the composite label for each of the eight unsupervised acoustic clusters obtained with the Leaf clustering method** (see Fig 12D).

## Data Availability

The Sound files from the Bora-Bora recording campaign, the configuration file (*CoralSoundExplorer* processing part input file) and generated data storage file (*CoralSoundExplorer* processing part output file) are archived at zenodo.org with the DOI 10.5281/zenodo.10794462. The python script used to generate the results of the section *IV. Detailed Methodology* and Fig 17, 18 & 19 are provided in the same zenodo.org deposit. Version 1.0.18 of CoralSoundExplorer, which was used for the work presented here, is available on the GitHub repository: https://github.com/sound-scape-explorer/coral-sound-explorer/archive/refs/tags/v1.0.18.zip.

## Acknowledgements

This research has been funded by the University of Saint-Etienne, the CNRS, the Inserm, Saint-Etienne Metropole, the Ecole Pratique des Hautes Etudes, the Institut Universitaire de France (NM and RE), and the Labex CeLyA. This publication was completed while NM was Visiting Miller Professor at the University of California, Berkeley (Miller Institute for Basic Research in Science).

## Author Contributions

**Name abbreviations:** LM: Lana Minier; JR: Jérémy Rouch, BS: Bamdad Sabbagh, FB: Frédéric Bertucci, EP: Eric Parmentier, DL: David Lecchini, FS: Frédéric Sèbe, NM: Nicolas Mathevon, RE: Rémi Emonet

**Conceptualization:** LM, JR, DL, FS, NM, RE.

**Bora-Bora Data curation:** LM

**Formal analysis:** JR, RE

**Funding acquisition:** DL, FS, NM, RE **Investigation:** LM, JR, DL, FS, NM, RE

**Methodology:** JR, BS, FS, NM, RE

**Project administration:** DL, FS, NM, RE

**Software:** JR, BS, FS, NM, RE

**Supervision:** DL, FS, NM, RE

**Validation:** All authors

**Writing - original draft:** LM, JR, DL, FS, NM, RE

**Writing - review & editing:** all authors.

